# An integrated stereotaxic injection architecture for mouse intracranial surgeries with AI-driven domain expertise

**DOI:** 10.64898/2026.09.21.753371

**Authors:** Mir Abbas Raza, Reilly Downing, Owen Beer, Megan LaBelle, Sanjay Madala, Kassidy Schroeder, Krishna Bhavithavya Kidambi, Aaron Sathyanesan

## Abstract

Stereotaxic intracranial microinjection in mice is a critical step in neuroscience workflows, enabled by robust hardware design, precise operation, and neuroanatomical expertise. Manual injections are prone to error and stereotax-mountable commercial automated injectors are extremely expensive. Further, the knowledge needed for performing accurate intracranial injections and surgeries is heterogeneous and disparate. Here we demonstrate the design and characterization of the UD Neuroinjector software-hardware-knowledge architecture, which serves to both reduce the cost of performing precisely controlled stereotaxic microinjections and provide a web platform for planning accurate targeting of specific mouse brain regions. We designed the UD Neuroinjector using low-cost hardware and 3D-printed parts, with a total build-cost of ∼$300 — less than 0.1× the cost of a commercial injector. Arduino-based firmware enables programmable flow rates, automated injection and extraction, and manual joystick control. Capillary-fluid measurements show stable, highly linear displacement (*R*^2^ > 0.93) at commonly used injection rates. In a head-to-head comparison with a commercial injector, *in vivo* DAPI injections into the mouse thalamus showed comparable spread and DAPI^+^ cell counts. For effective experimental design of intracranial injections, the web-based components of our architecture provide AI-enabled assistance and atlas-guided navigation. We used a hybrid retrieval-augmented generation (RAG) approach to ground AI-assisted protocol guidance for mouse intracranial surgeries using a corpus of 3,738 papers. Using the Common Coordinate Framework version 3 (CCFv3), we designed an interactive tool for identifying target coordinates. Together, the UD Neuroinjector architecture integrates and enables efficient knowledge discovery, experimental design, planning, and implementation of mouse intracranial surgeries.

## 1 Introduction

Stereotaxic microinjections into the rodent brain are a fundamental part of a wide range of neuroscience experiments. The efficiency of approaches such as neuroanatomical tracing, induced cell ablation studies, and viral induction of biosensors for *in vivo* physiology or optogenetics largely depend on stable injection of chemicals or biologics at precise flow rates into the brain [1, 2]. While manual stereotaxic injections into the rodent brain are still commonly practiced, neuroscience research labs often resort to using commercial automated microinjection systems. These commercial injector systems are very expensive (upwards of $3000) and can be cost-prohibitive in resource-poor contexts [3, 4]. However, few open-access and low-cost solutions for automated stereotaxic microinjectors exist. Importantly, head-to-head *in vivo* validation in comparison to commercial injectors is lacking.

In addition to the automation of intracranial injections, researchers performing these injections also rely on knowledge components such as mouse or rat brain atlases and previously published protocols [5, 6]. Given the sheer anatomical and cellular complexity combined with the small size of the mouse brain, the precision of injection into a particular brain region depends on the accurate identification of stereotaxic coordinates of the said region. While few recent tools have been designed for planning viral injections, these are most appropriate for planning neural recording probe trajectories which require additional considerations of pitch and yaw angles in 3D space [7, 8]. For stereotaxic viral injections in general, the first main consideration includes the identification of target coordinates along the antero-posterior (AP), mediolateral (ML), and dorsoventral (DV) axes using a standardized coronal brain atlas. The second main consideration includes prior knowledge of successful stereotaxic targeting from peer-reviewed publications. However, no tool has incorporated the rich corpus of stereotaxic neurosurgical information from previously published studies in the mouse. We designed an end-to-end open-source architecture – the UD Neuroinjector Open Source Ecosystem (OSE) – that includes hardware, software, and knowledge components that can be applied across the entire workflow of stereotaxic intracranial injection into the mouse brain. We built the UD Neuroinjector using fused filament fabrication

3D printed parts, low-cost electronic and mechanical components, and simple serial control via an Arduino microcontroller board. The UD Neuroinjector is capable of stable flow rates that are commonly used in mouse intracranial injection protocols. *In vivo* intracranial DAPI injections showed that the UD Neuroinjector is similar in performance to that of a commercial injector. Finally, we used a hybrid retrieval augmented generation (RAG) approach to build domain expertise in mouse brain surgery. This information is easily accessible through an LLM via a chat-based interface. Using the Allen Brain Atlas Common Coordinate Framework version 3 (CCFv3), we designed a simple web app to complement domain expertise through AI-assisted protocol planning. Our system enables comprehensive workflow planning, troubleshooting, and implementation of successful stereotaxic intracranial injections into the mouse brain, which is a cornerstone technique in modern neuroscience.

## 2 Results

### 2.1 Hardware and software layer

#### 2.1.1 UD Neuroinjector design and control

We used off-the-shelf electronic and mechanical components combined with 3D-printed mounting and housing for the UD Neuroinjector, with a net cost of 289 USD (Fig. 1a-b). We used a NEMA 8 bipolar stepper motor (1.8^*°*^ full step angle; 200 full steps per revolution, i.e., 360^*°*^) coupled to an ultraprecision ⅛ inch lead screw (0.6096 mm rev^*−*1^) to drive a flange nut connected to a 3D-printed plunger depresser that slides along a linear guide rail (Fig. 1b).

**Fig. 1.**
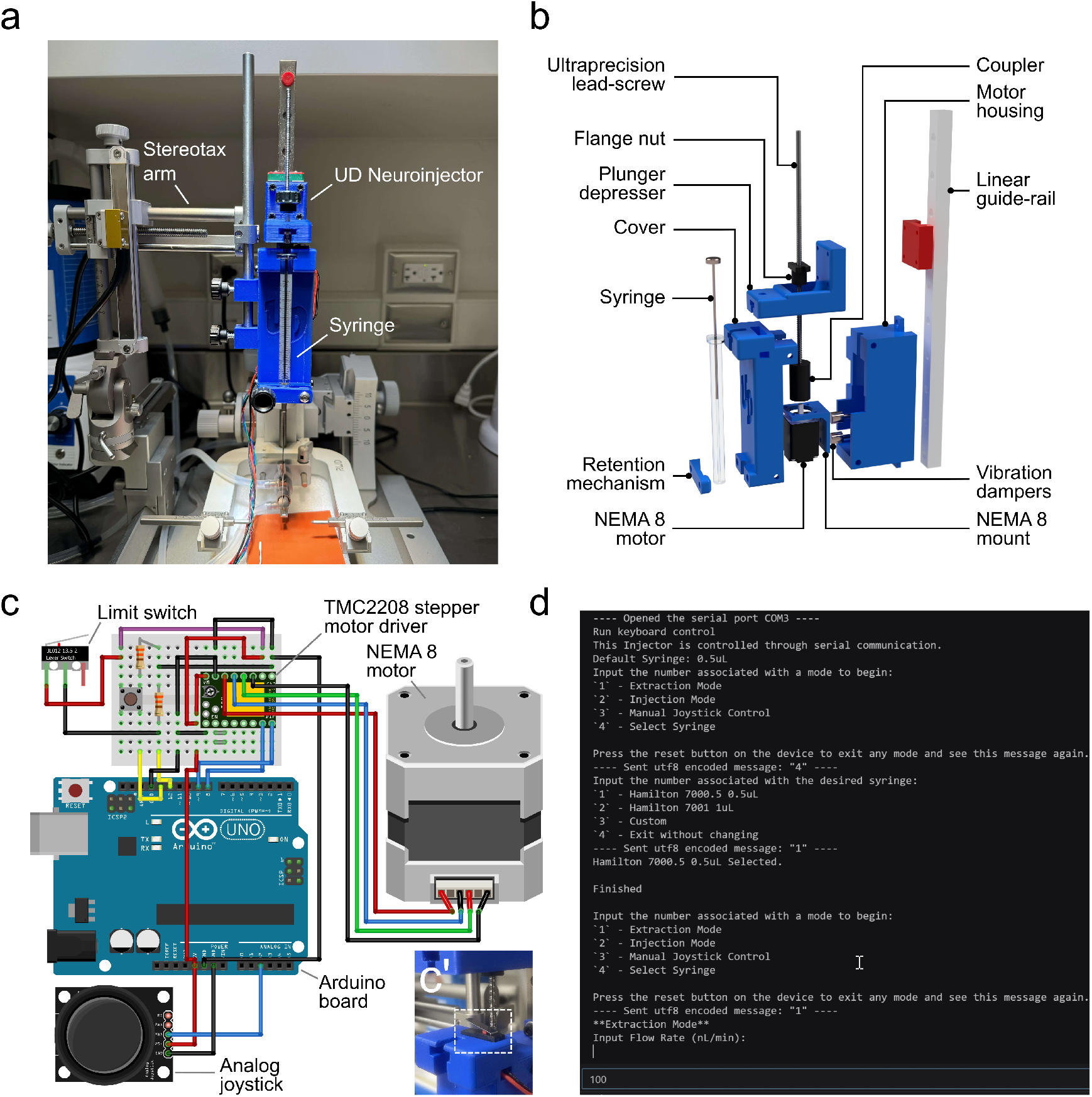
UD Neuroinjector Hardware Design and Control. **a**. A photograph of the UD Neuroinjector mounted onto the arm of a mouse stereotaxic setup. **b**. Exploded view of the components of the UD Neuroinjector **c**. Wiring diagram (fritzing) of Arduino microcontroller-based control of the NEMA 8 motor in the UD Neuroinjector; inset (**c’**) shows limit switch placement to prevent damage.**d**. Serial-programming-based interface for controlling the UD Neuroinjector. The script for UD Neuroinjector control and serial programming can be downloaded from the UD Neuroinjector GitHub repository.

A 3D-printed NEMA 8 mount holds the motor in place within the motor housing with vibration dampers for stable operation. The outer part of the cover for the motor housing has a recess to hold the upper lip of the syringe barrel in place, with the lower part of the barrel secured to the cover using a retention mechanism (Fig. 1b). This helps hold the syringe securely in place during injections. The plunger depresser recess is designed to hold the plunger button in place so it can move precisely along the vertical axis. The side of the motor housing facing the stereotaxic arm has two locations along which the standard-size stereotaxic arm adapter can be passed through to mount the UD Neuroinjector (Fig. 1a). A pair of set screws secure the injector to the stereotaxic arm adaptor.

The NEMA 8 stepper motor is driven by a TMC2208 stepper motor driver (Fig. 1c), which is configured for microstepping yielding 1600 microsteps per revolution. Given this mechanical configuration, the theoretical displacement resolution is calculated as follows:

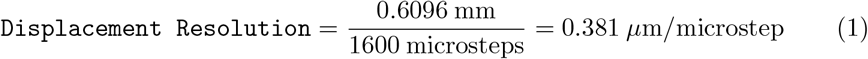

An Arduino UNO board controls the NEMA 8 motor and TMC2208 stepper motor driver (Fig. 1c). An analog joystick controls manual injector up/down movement. A limit switch (Fig. 1c) checks if the plunger has been fully depressed and, if switched ON, prevents further stepping. A simple, interactive, serial programming interface controls the injector (Fig. 1d).

#### 2.1.2 Flow rate validation for UD Neuroinjector

Since intracranial injections into the mouse brain depend on stable and relatively low flow rates, we sought to determine how stably the UD Neuroinjector can dispense a defined volume (150 nl) at different flow rates commonly used in mouse intracranial injections. Using a high-resolution macro video camera, we recorded water meniscus displacement in a 50 *µ*l capillary tube to measure injection flow rate stability (Fig. 2a). Since water is injected into the capillary, the meniscus is displaced upward towards a black band (reference) marked on the capillary tube.

**Fig. 2.**
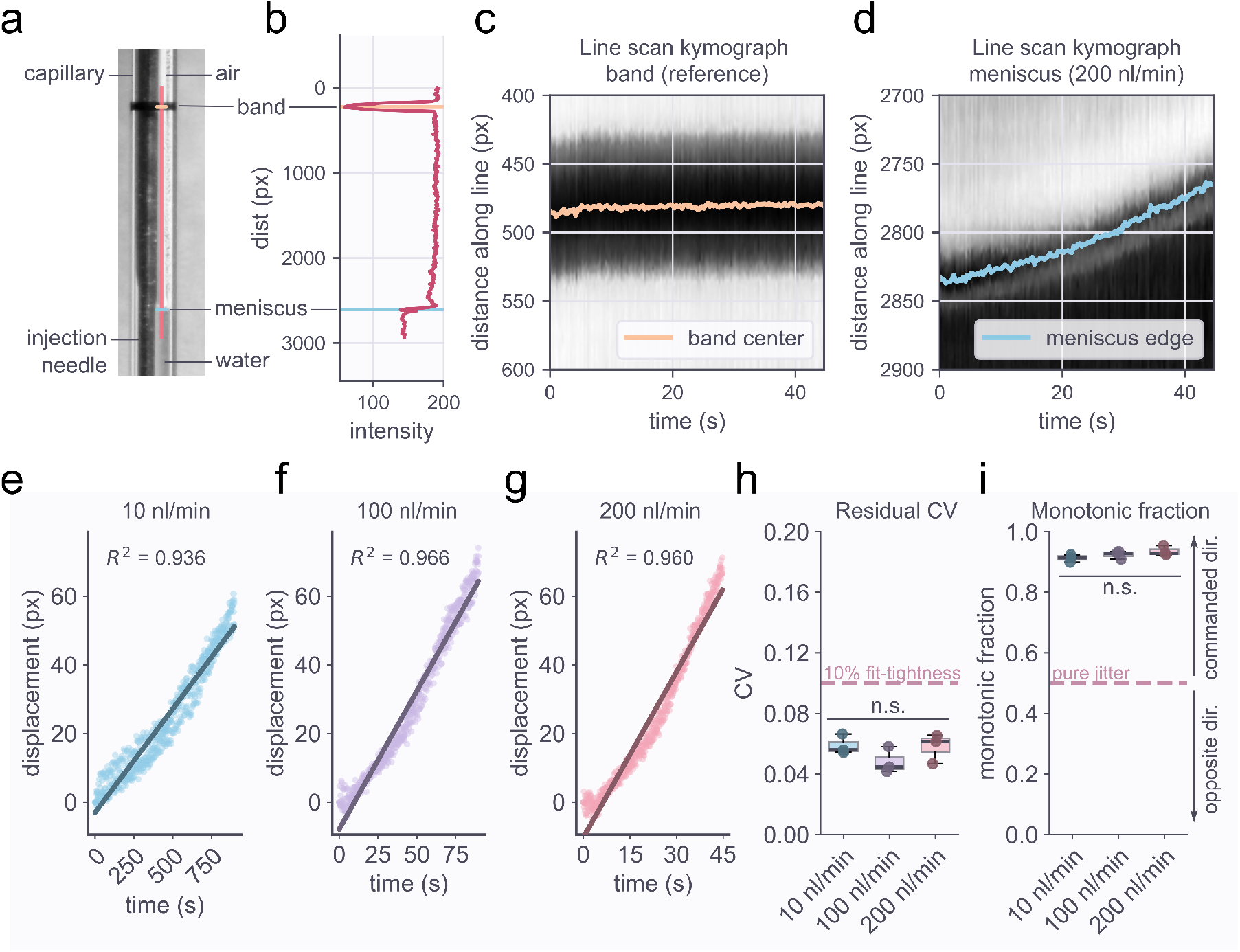
Flow rate validation for UD Neuroinjector. **a**. Fluid capillary displacement setup showing the black band (‘band’) reference, syringe injection needle, and the water meniscus at the air-water interface inside the capillary. **b**. Line intensity scan profile of the orange line in panel a showing the inverted peak (orange) corresponding to the reference black band as well as the meniscus profile peak (blue). **c**. Line scan kymographs for the center of the black reference band (orange) and **d**. the meniscus edge (blue) for a representative 200 nl/min injection. **e**. Meniscus relative displacement over the course of the injection for 10 nl/min **f**. 100 nl/min and **g**. 200 nl/min flow rate. **h**. Residual CV comparison between the different flow rate measurements (no statistically significant difference [n.s.] between CV across flow rates; N = 3 injections per group; One-way ANOVA, P = 0.3043). **i**. Comparison of monotonic fraction across different flow rates (no statistically significant difference [n.s.] between monotonic fraction across flow rates; N = 3 injections per group; One-way ANOVA, P = 0.2071)

Using automated line intensity scan analysis, we tracked the movement of the meniscus (Fig. 2a–b, blue line) with respect to the reference band (Fig. 2a–b, orange line). Representative line scan kymographs show no noticeable drift in the position of the black reference band (Fig. 2c) and a steady upward movement of the meniscus edge (Fig. 2d).

The UD Neuroinjector showed stable linear meniscus displacement at injection rates of 10 nl/min (Fig. 2e; *R*^2^ = 0.936), 100 nL min^*−*1^ (Fig. 2f; *R*^2^ = 0.966), and 200 nl/min (Fig. 2g; *R*^2^ = 0.960). Fit tightness using residual CV did not show significant statistical differences across different flow rates (Fig. 2h).

We also measured monotonic fraction as a measure of fractional forward progression over a rolling window. Mean monotonic fraction exceeded 0.9 for all injection flow rates tested (Fig. 2i). We also did not notice any statistically significant difference in monotonic fraction across flow rates, indicating stable forward injection across conditions. For every injection, the plunger moved completely to the bottom of the barrel, indicating that the entire 150 nl had been dispensed.

#### 2.1.3 In vivo head-to-head performance of the UD Neuroinjector

To perform head-to-head comparison of the UD Neuroinjector with a commercial stereotax-mountable injector (Quintessential Stereotaxic Injector, Stoelting Co., Wood Dale, IL, United States), we injected the DNA-intercalating fluorescent stain 4^*′*^,6-diamidino-2-phenylindole (DAPI) into the mouse lateral posterior thalamic nucleus (AP *∼* −1.2 to −1.4 mm, ML*∼* 0.9 to 1.2 mm, DV*∼* − 2.6 to −2.8 mm) just ventral to the medial dentate gyrus (Fig. 3a)

**Fig. 3.**
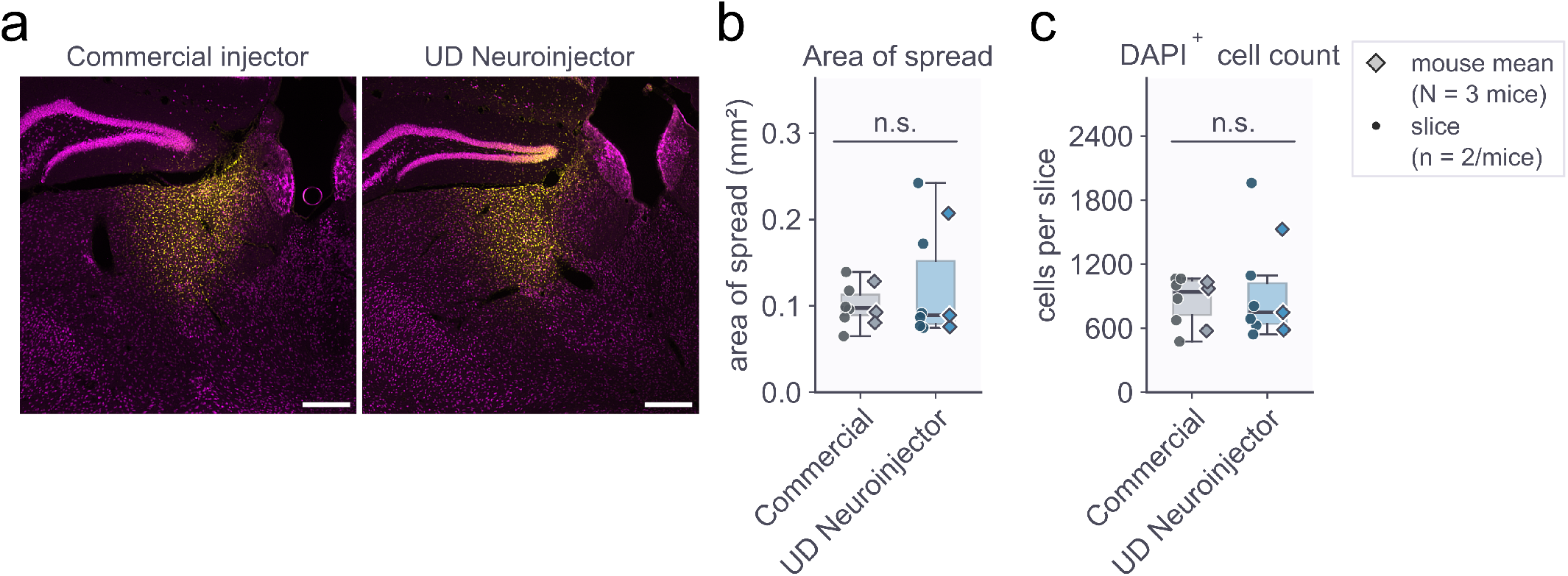
*In vivo* head-to-head performance of UD Neuroinjector vs. a commercial injector. Representative fluorescence microscopy images of mouse brain tissue sections following stereotaxic surgery and injection of 100 nl of 5 mg/ml DAPI solution using the Quintessential Stereotaxic Injector (QSI, Stoelting Company) (left image) and the UD Neuroinjector (right image) (AP ∼*−*1.2 to *−*1.4 mm, ML*∼* 0.9 to 1.2 mm, DV*∼ −*2.6 to *−*2.8 mm); DAPI shown in yellow, anti-NeuN pan-neuronal marker shown in magenta. **b**. Area of spread comparison between the commercial injector and the UD Neuroinjector shows no statistically significant (n.s.) differences in area of spread (N = 3 mice, n = 2 slices/mouse; Welch’s independent-samples t-test on per-animal mean measurement; Welch’s t = -0.5313, two-sided P = 0.6392). **c**. No statistically significant differences in DAPI+ cell counts (N = 3 mice, n = 2 slices/mouse; Welch’s independent-samples t-test; Welch’s t = -0.2889, two-sided P = 0.792) between the commercial injector and the UD Neuroinjector. Scale bar for panel a = 150 µm.

Computational image analysis of DAPI staining showed that the UD Neuroinjector performed comparably to the commercial injector both in terms of the total area of spread across the injected region (Fig. 3b; Welch’s t-test on per-mouse measurements; no statistically significant difference between groups; *N* = 3 mice per group; Welch’s t = -0.53; df = 2.47; P = 0.64) as well as the automated DAPI^+^ cell counts (Fig. 3c; Welch’s t-test on per-mouse measurements; no statistically significant difference between groups; *N* = 3 mice per group; Welch’s t = -0.29; df = 2.92; P = 0.79). The UD Neuroinjector thus has similar performance as a standard automated commercial injector in terms of *in vivo* intracranial injections.

#### 2.1.4 Routine laboratory use

In everyday practice, we have used the UD Neuroinjector to inject adeno-associated viral (AAV) vectors into the thalamus and cerebellum at multiple stereotaxic coordinates. A retrospective analysis of histologically confirmed tissue data in our lab between September 1, 2025, and May 12, 2026, showed that the UD Neuroinjector had a success rate of 8/10 injections, similar to the commercial injector with a success rate of 9/10 injections that showed intended expression in the target brain region. In cases of the failed injections for both injectors we could not accurately determine the cause of failure in terms of histological expression.

### 2.2 Domain-expertise and knowledge layer

Stereotaxic injections into the mouse brain are challenging at both the hardware-software level as well as the knowledge and domain-expertise level. Having provided a stable and robust open-source solution using the UD Neuroinjector described above, we proceeded to design the knowledge and domain-expertise component necessary for successful intracranial injections into the mouse brain (Fig. 4).

**Fig. 4.**
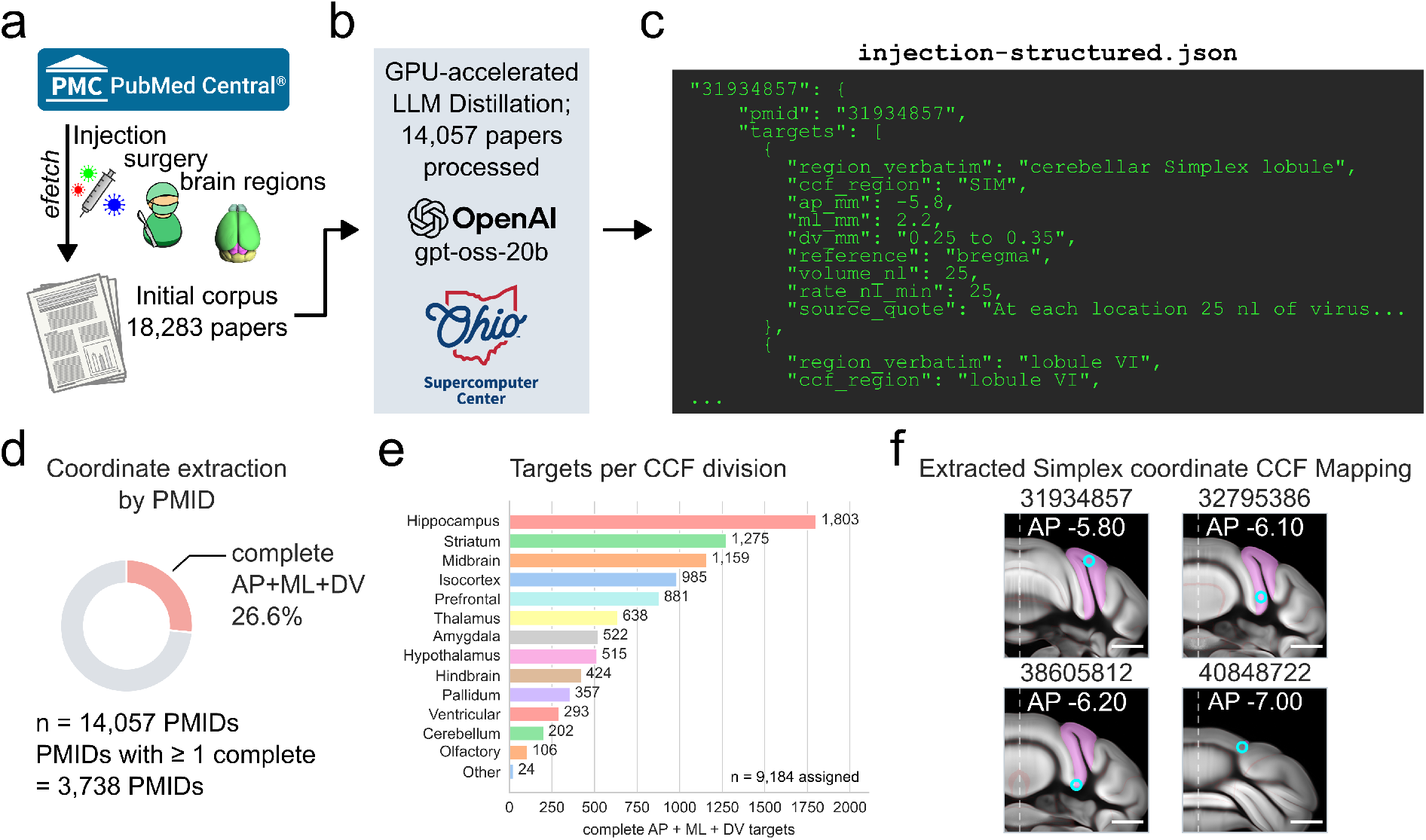
Corpus-grounded AI-driven domain expertise and knowledge layer for the UD Neuroinjector ecosystem. **a**. Initial scoping of PubMed Central (PMC) for search terms associated with injection, surgery, and mouse brain regions yielded an initial corpus of 18,283 papers. We used GPU-accelerated distillation of the papers using the `gpt-oss-20b` open source model on the Ohio Supercomputer and obtained **c**. a structured file, `injection-structured.json.` **d**. The `injection-structured.json` file contains 3,738 PMIDs with at least one complete (antero-posterior – AP, mediolateral – ML, and dorsoventral – DV) coordinate set representing 26.6% of the 14,057 PMIDs. **e**. An analysis of the targets per common coordinate framework (CCF) division (n = 9,184 targets assigned) reveals the spread of the coordinates, with the hippocampus having the largest number of previously reported coordinates followed by the striatum. **f**. Proof-of-feasibility mapping of extracted coordinates (cyan circles) onto the CCFv3 coordinate system for the cerebellar simplex (magenta) found in PMIDs 31934857 (AP -5.80 relative to bregma), 32795386 (AP -6.10 relative to bregma), 38605812 (AP -6.20; relative to bregma but rederived from lambda – 4.20), and 40848722 (AP -7.00; relative to bregma but rederived from lambda – 4.20).

#### 2.2.1 Initial corpus scoping

We developed a Node.js/JavaScript trawler pipeline `(ingest-protocols.js)`that queried the NCBI PubMed and PMC databases using the NCBI E-utilities `efetch` endpoint [9] for mouse stereotaxic injections into brain regions. This broad-based approach ensured that we would get maximal coverage of the available corpus, retrieving article metadata and available full text. When `efetch` was unable to extract full text, we used the PMC BioC API [10] as a fallback.

We focused on passages that included the results, text from the figure captions and legends, and the methods section. Following deduplication, this text was assembled into the `surgical-protocols.json` file. This JSON file has stereotaxic-surgery-related text for a total of 18,283 papers (Fig. 4a).

#### 2.2.2 LLM-based corpus distillation

To sort through these large passages of text and identify mouse brain stereotaxic coordinates, we used an LLM-based approach on the Ohio Supercomputer (Fig. 4b). First, we used keyword-based filtering to identify papers with mouse, stereotaxic, coordinate, and injection-related signals and build vLLM batch prompt files. We then used a distillation SLURM script on OSC to run OpenAI’s open-weight model `gpt-oss-20b` (21B parameters) [11] over the prompts.

The final outputs were collected into the `injection-structured.json` file (Fig. 4c), which contains coordinates (at least one type—AP, ML, or DV—extracted) for 14,057 unique PMIDs, of which 26.6% (3,738 PMIDs) have at least one complete set (AP, ML, and DV) (Fig. 4d). A total of 13,044 target regions were extracted, from which 9,184 were confidently assigned to a particular CCF division (Fig. 4e), with the hippocampus and striatum comprising a large percentage of the targets. We independently curated a ground-truth dataset (Online Resource 1) for targets previously reported in the Nucleus Accumbens (Acb), Hippocampal CA1 region (CA1), and the Caudate putamen (CP). We used this dataset to test the accuracy of the LLM-extracted coordinates (Table 1). In comparison to ground-truth, the average accuracy of LLM-distilled stereotaxic coordinates by brain region was 97.7% (Acb: 100%; CA1: 93.1%; CP: 100%). The average accuracy by stereotaxic coordinate axis was 99.2% (AP: 98.8%; ML: 100%; DV: 98.8%).

**Table 1.**
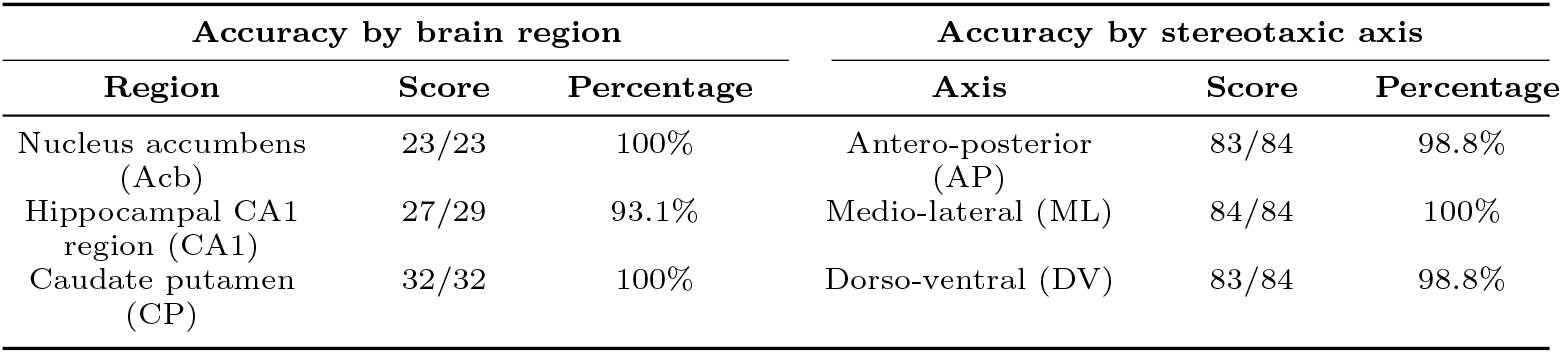
Accuracy of LLM-distilled coordinates by brain region and by stereotaxic axis.

#### 2.2.3 Identification of mouse brain stereotaxic coordinates using RAG

We implemented a simple UD Neuroinjector Assistant chat module that uses deterministic retrieval to identify stereotaxic coordinates grounded in the scientific corpus (Online Resource 2) [12, 13]. The UD Neuroinjector Assistant uses a React client to load the structured coordinates and relevant extracted passages. The system tokenizes user input and ranks candidate targets and literature records based on lexical similarity, anatomical-region matching, PMID match, and coordinate-target relevance. Regional queries are also given a “structure-aware” score.

To ensure maximal fidelity to mouse-specific stereotaxic coordinates, a mouse-specific filter was enforced. Highest-scoring candidates receive priority in the output response, which contains the PMID, anatomical region as stated or CCFv3-matched, AP/ML/DV coordinates, injection volume, flow rate, and the original source quote. Strict grounding rules are enforced for the language model (GPT 5.5 or GPT 5.4) to only cite papers found in `injection-structured.json.`

Additionally, to close the hardware-software-knowledge loop, we also made hardware troubleshooting and syringe information relevant to the UD Neuroinjector available to the assistant. The UD Neuroinjector Assistant thus serves as a “one-stop-shop” for stereotaxic target planning using a RAG-based approach with deterministic lexical and anatomical-context matching.

#### 2.2.4 A simple CCFv3-based atlas module for target coordinate planning

We designed a simple and interactive atlas module using the Allen Mouse Common Coordinate Framework version 3 (CCFv3) for effective stereotaxic target planning [12]. The module uses the CCFv3 and the Allen structure ontology to map voxels onto anatomical structures and region names. We use coronal slices indexed by AP axis position and overlaid with anatomical boundaries. Users can search for a particular location using a dropdown or click-to-target a particular brain region.

For a selected location on a brain region within a defined AP slice, ML and DV coordinates are calculated and displayed in millimeters relative to the CCFv3 reference axes. Targets can also be resolved manually by users. Once a target with AP, ML, and DV (w.r.t. Bregma) coordinates has been identified, this can be queried by the LLM to check for consistency with previously published papers, as seen in the screen-recording (Online Resource 3).

Using the atlas in combination with corpus-grounded AI-driven domain expertise may serve as an effective strategy for optimal stereotaxic targeting. We verified the accuracy of extracted coordinates for the cerebellar simplex (or simple lobule) from specific papers (unique PMIDs) by mapping them onto the closest-matching AP slice images in our atlas module reference system (Fig. 4f). We observed striking concordance for the extracted coordinates (Fig. 4f, cyan circle) within the simplex (Fig. 4f, magenta shaded region).

## 3 Materials and Methods

### 3.1 Animals

All animal procedures were performed according to the Institutional Animal Care and Use Committee of the University of Dayton (Protocol 023-04) and the Guide for the Care and Use of Laboratory Animals (NIH). We used adult C57BL/6 and B6EiC3Sn.BLiAF1/J mice in injection experiments. Both male and female sexes were used for injection experiments.

### 3.2 Surgery and stereotaxic intracranial injections

Mice were weighed and subjected to anesthesia induction (3% Isoflurane) with isoflurane in a chamber linked to a table-top isoflurane system (RWD Life Science Inc., Dover, DE, United States). After induction, mice were placed on a temperature-controlled heating stage (ALA Scientific, Farmingdale, NY, United States). Body temperature was monitored using a rectal probe connected to the heating stage to maintain optimal body temperature. The snout was then positioned inside an anesthesia mask (RWD Life Science Inc., Dover, DE, United States) and 3% Isoflurane was supplied through this line.

The mouse was then positioned carefully in the digital stereotaxic frame (RWD Life Science Inc., Dover, DE, United States) by placing the ear bars slowly into the ear canal. Lubricant ointment (Paralube vet ointment, Dechra, Boston, MA, United States) was topically applied to the eyes to prevent drying. Once the breathing rate stabilized (1–1.5 breaths per second), isoflurane was reduced to *∼* 1.5% – 2.5% for anesthesia maintenance.

The fur above the skull was closely clipped using surgical scissors. Betadine was applied to the skin above the skull using a sterile cotton swab. 0.5% Lidocaine (2.5–4 mg/kg; Vedco, St. Joseph, MO, United States) was injected locally in the skin above the skull as a local anesthetic. A single incision was made along the midline to expose the skull using a disposable scalpel (Foster City, CA, United States). Bleeding if any was stemmed using Sugi^®^ Sponge Points (Kettenbach, Huntington Beach, CA, United States). Sterile cotton swabs were used to clear away any tissue above the skull.

Once the skull was clearly visible, 3% hydrogen peroxide was applied to the skull to visualize the bregma. Saline was then immediately applied to neutralize the hydrogen peroxide. Excess solution was cleared away using sterile cotton swabs. The bregma was then recorded digitally, following which a 0.6 mm burr hole was drilled in the skull at AP∼ −1.2 to −1.4 mm, ML ∼0.9 to 1.2 mm, using an electric dental drill. Sterile saline was applied to the holes to prevent drying. The dura was carefully removed using fine forceps. 100 nl of 5 mg/ml DAPI solution (10.9 mM solution reconstituted in saline; BioLegend, San Diego, CA, United States) was injected into the thalamus at a depth (DV) of ∼−2.6 to −2.8 mm using a 1 *µ*l syringe (7001KH, PN: 80100, Hamilton Company, Reno, NV, United States) mounted on an automated injector (either the UD Neuroinjector or the Quintessential Stereotaxic Injector, Stoelting Co., Wood Dale, IL, United States) at a flow rate of 100 nl/min.

Two minutes after the DAPI solution had been completely injected into the site, the syringe was gently retracted. Vetbond tissue glue (3M, St. Paul, MN, United States) was applied to glue the skin flaps back. Meloxicam (5 mg/kg) was injected into the scruff for analgesia, and isoflurane was gradually reduced. The mouse was allowed to rest on the stage till it regained consciousness and generated active movement. It was then transferred to its home cage and monitored for 1 hour following completion of surgery. The mouse was monitored once every 12 hours for signs of pain, dehydration, reduction of activity and discomfort. The mouse was then transcardially perfused within 24 hours after surgery, and the brain was used for tissue processing and immunohistochemistry. We performed one injection per mouse into one hemisphere.

### 3.3 Tissue processing and immunohistochemistry

Mice injected with DAPI were anesthetized with isoflurane, and following tailpinch-verification, transcardially perfused with ice-cold 0.1 M Phosphate Buffered Saline (PBS) followed by 4% paraformaldehyde (PFA). The brain was dissected out of the skull and post-fixed in 4% PFA with 30% Sucrose overnight at 4 ^*°*^C after which the brain was stored in 0.1 M PBS with 30% sucrose at 4 ^*°*^C till the tissue processing stage. The brain was cut into free-floating 35 *µ*m coronal sections using a freezing-stage (BFS-40MPA, Physitemp Instruments, Clifton, NJ, United States) mounted on a sliding microtome (HM 430, Epredia, Kalamazoo, MI, United States). Sections were labeled using standard immunohistochemical procedures.

To visualize neurons in DAPI-stained sections, we used an anti-NeuN primary antibody (1:2000; Cat. No. 266 004, Synaptic Systems GmbH, Goettingen, Germany) followed by an Alexa Fluor 488 anti-guinea-pig secondary antibody (Code: 706-545-148, Jackson Immunoresearch Laboratories, West Grove, PA, United States). The tissue was then mounted onto Fisherbrand Superfrost Plus Microscope slides (Fisher Scientific Company, Hanover Park, IL, United States), with a few drops of Fluoromount-G (SouthernBiotech, Birmingham, AL, United States) and covered with a cover glass (Fisher Scientific Company, Hanover Park, IL, United States).

### 3.4 Fluorescence microscopy and imaging

We used an Olympus BX53 epifluorescence microscope equipped with an LED illumination system, DAPI/FITC/Cy3/Cy5 filters, an Olympus DP23M monochrome camera, and Olympus CellSens software (Evident Scientific Inc., Needham, MA, United States) for fluorescence imaging of DAPI^+^ and NeuN^+^ neurons. Images were captured using a 4*×* Olympus air objective in the TIFF format using the CellSens software at 3088*×*2076 pixel resolution.

### 3.5 Automated cell quantification

We used ImageJ Fiji [14] and the pixel classification tool Ilastik [15] for automated DAPI^+^ cell quantification. We converted the TIFF microscopy images to PNG and used these files as input training data to train an Ilastik segmentation model. Ilastik simple segmentation files were then analyzed on Fiji using simple thresholding. For DAPI^+^ cell counting we used the ‘Analyze particles’ tool. For calculating area of spread we first applied a gaussian blur followed by thresholding to binarize the area with DAPI^+^ cells and the rest of the brain section area with no DAPI. After this we used the ‘Analyze particles’ tool to quantify the area of spread in mm^2^.

### 3.6 Fluid displacement video acquisition and analysis

We inserted a Hamilton syringe injection needle (Hamilton 7000.5, PN: 86250, Hamilton Company, Reno, NV, United States) into a 50 µL glass capillary tube (Yankee Disposable Micropet, BD, Parsippany, NJ, United States) pre-filled with distilled water such that there is little air space between the black reference band on the capillary and the water meniscus. The bottom of the capillary was sealed off with parafilm. The capillary was held upright using a clamp. We used a Samsung Galaxy S25 Ultra (Sony IMX854 camera sensor) mounted in a secure holder for acquiring ultrahigh definition video at 30 frames s^*−*1^ and 25*×* magnification for a pixel resolution of 3840 *×*2160.

For analysis, we used features on frame-by-frame line intensity scans that covered both the black reference band as well as the meniscus (similar to Fig. 2a–b). We first sought to reduce the effect of camera jitter by calculating the vertical axis coordinate of the meniscus, *y*_meniscus_(*t*), relative to the black band reference on the capillary, *y*_band_(*t*), for *N ≈* 200 sampled data points per injection:

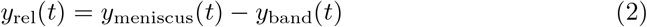

We fitted displacement over time data using an ordinary least squares linear model.

We then calculated the residual coefficient of variation *CV* :

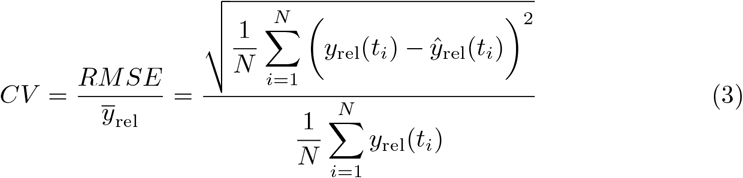

Where *ŷ*_rel_(*t*_*i*_) is linear regression fit of *y*_rel_ at time *t*_*i*_. We also calculated monotonic fraction, *f*_mono_, as:

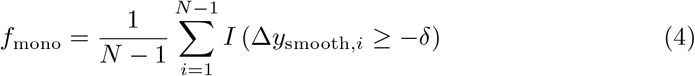

Where:

- *y*_smooth_(*t*_*i*_) = median (*y*_rel_(*t*_*i−*3_), …, *y*_rel_(*t*_*i*+3_))
- Δ*y*_smooth,*i*_ = *y*_smooth_(*t*_*i*+1_) *− y*_smooth_(*t*_*i*_)
- *δ* = 0.02 *×* median (|Δ*y*_smooth,*i*_|)
- *I*(*·*) = 1 if true, 0 otherwise.

### 3.7 UD Neuroinjector 3-D printing and assembly

The UD Neuroinjector assembly needs low-cost off-the-shelf components and 3Dprinted parts. We used a standard FFF (or FDM) printer to 3D-print the parts for the UD Neuroinjector. We have provided detailed documentation for step-by-step assembly of the UD Neuroinjector on the UD Neuroinjector OSE GitHub Repository [13]. The time needed to build the UD Neuroinjector once all the parts have been assembled is *∼* 30 minutes. We have also included all source STL files for 3D-printed parts as well as a bill of materials.

The default motor housing design helps secure the UD Neuroinjector onto the stereotaxic arm on the left-hand side of the stereotaxic frame. However, in the STL folder of the GitHub repository we have also provided a file `(Motor Housing righthanded.STL)` that can be used to 3D-print a motor housing that can be secured to a “right” stereotaxic arm if needed.

### 3.8 Arduino-based actuation

We used an Arduino UNO R3 microcontroller board with a prototype expansion shield for controlling the UD Neuroinjector. A NEMA 8 bipolar motor (1.8^*°*^ step angle, 28 mm body length, 0.2 A rated current, 1.6 N cm holding torque; Amazon, Seattle, WA, United States) coupled to an ultra-precision 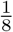 -inch lead screw (41.7 threads in^*−*1^, 0.6096 mm pitch; McMaster-Carr, Elmhurst, IL, United States) via an aluminum rigid shaft coupler (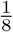-inch-to-5 mm; Amazon, Seattle, WA, United States) provided linear actuation.

To increase stability and smoothness of operation, the plunger depresser is mounted onto an MGN12H carriage block that slides along a 250 mm linear guide rail (Amazon, Seattle, WA, United States). The full steps per revolution, *N*_step_, of the NEMA 8 are calculated as:

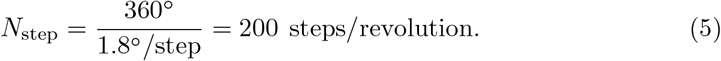

A TMC2208 stepper motor driver (Amazon, Seattle, WA, United States) enables microstepping. Microsteps per revolution, *N*_micro_, and degrees per microstep, *θ*_micro_, are calculated as:

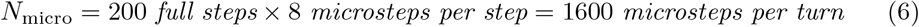

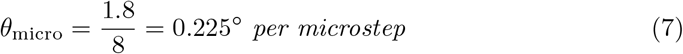

Linear actuation per turn, *Actuation*_turn_, i.e. for 360^*°*^, considering lead screw dimensions (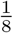 -inch) is 0.6096 mm. Linear actuation for step (*Actuation*_step_), and microstep (*Actuation*_micro_), is calculated as:

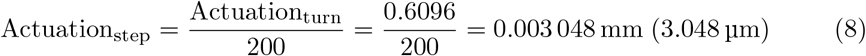

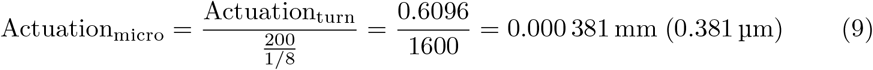

### 3.9 Firmware and serial programming

The Arduino control script `(injector control.ino)` is built on the open-source Accel-Stepper library [16], driving the TMC2208 through a step/dir interface (digital pins 8 and 9) over a 115200-baud serial link. The firmware implements four operating modes: extraction, injection, manual joystick control, and syringe selection, chosen by serial commands. Motion is executed as relative displacement from a reference position that is re-zeroed at the start of each mode. Syringe selection configures the plunger gradation *g*, defined by the firmware variable `syringe div`, in nanoliters of fluid displaced per millimeter of plunger travel. Compile-time constants define two presets: `HAM 7000 5 DIV =` 25*/*3 *≈* 8.33 nL mm^*−*1^ for the Hamilton 7000.5 (0.5 µL; PN: 86250, Hamilton Company, Reno, NV, United States) syringe and `HAM 7001 DIV =` 50*/*3 16.67 nL mm^*−*1^ for the Hamilton 7001 (1 µL; PN: 80100, Hamilton Company, Reno, NV, United States) syringe. Selecting a syringe (mode 4) copies the corresponding constant into `syringe div;` a custom gradation (1 nL mm^*−*1^ to 1000 nL mm^*−*1^) can be entered for any other syringe. The presets correspond to the full syringe volume delivered over the 60 mm plunger travel of the Hamilton 700-series microliter syringes (e.g., 8.33 nL mm^*−*1^ *×*60 mm = 500 nL for the 0.5 µL syringe).

For a commanded injection or extraction volume *V* (nL), the required plunger displacement is converted to microsteps as:

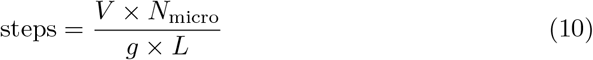

where *L* is the lead screw pitch (firmware variable lead, 0.6096 mm revolution^*−*1^) and *N*_micro_ = 1600 microsteps revolution^*−*1^ (firmware constant stepsprev). For a requested flow rate *F* (nL min^*−*1^), the firmware computes the required microstep rate, *S*, as:

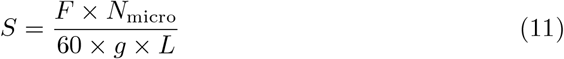

In the firmware code, Equations 10 and 11 are evaluated as `steps = vol * stepsprev / (syringe div * lead) and steppsec = (flow / syringe div) / lead / 60 * stepsprev,` respectively. The firmware validates all user input before motion begins, accepting flow rates of 0.1 nL min^*−*1^ to 1000 nL min^*−*1^ and volumes of 0.1 nL to 500 nL; the upper volume bound corresponds to the full stroke of the 0.5 µL syringe. For example, a 100 nL injection at 100 nL min^*−*1^ using the 0.5 µL syringe corresponds to 31,496 microsteps (12.0 mm of plunger travel) executed at 524.9 microsteps s^*−*1^, completed in 60 s. The table below (Table 2) maps the notation to its firmware representation.

**Table 2.**
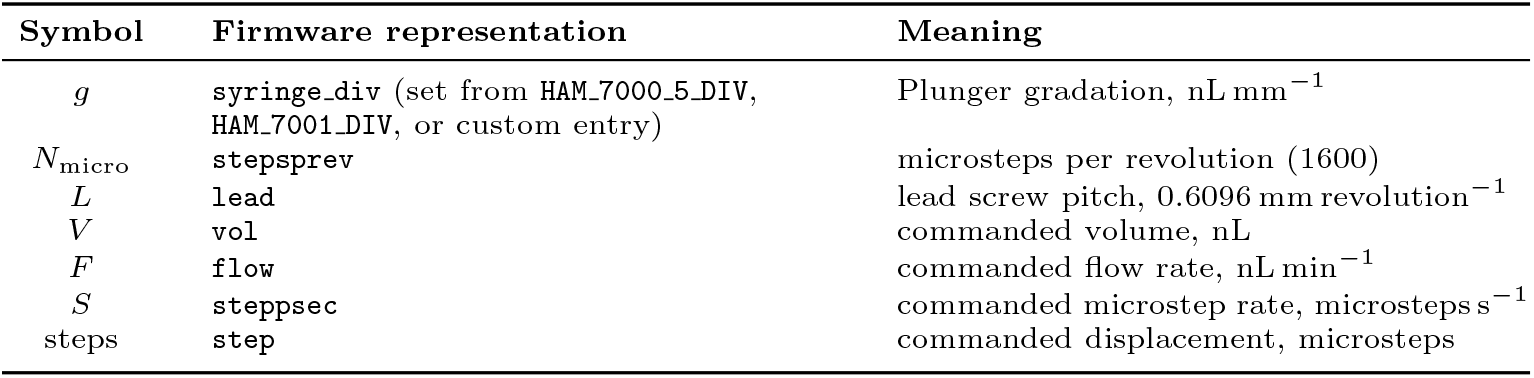
Mapping of mathematical notation to firmware representation.

### 3.10 Typical operation

Plunger displacement per microstep determines volumetric resolution of the UD Neuroinjector:

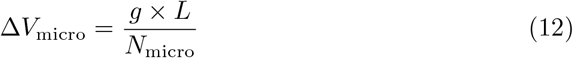

Where Δ*V*_micro_ is the volume delivered per microstep, which for a Hamilton 7000.5 (0.5 µL) syringe is *≈*0.0032 nL (3.2 pL) per microstep and *≈*0.0064 nL (6.4 pL) for the Hamilton 7001 (1 µL) syringe. The achievable flow-rate range is determined by the maximum step rate (8000 microsteps s^*−*1^). The accepted input range of 0.1 nL min^*−*1^ to 1000 nL min^*−*1^ maps to 0.53 microsteps s^*−*1^ to 5249 microsteps s^*−*1^ on the 0.5 µL syringe (0.26 microsteps s^*−*1^ to 2625 microsteps s^*−*1^ on the 1 µL syringe). The control script also reports percentage injection progress at 1 Hz.

In the manual joystick mode, Analog PIN A2 translates up-or-down (X) displacement from the joystick into a 600 nL min^*−*1^ plunger stroke in the corresponding direction for fast control. The downward plunger stroke is gated by the limit switch (PIN 12).

Follow the instructions below for a standard injection:

1. Mount the injector onto the stereotaxic arm. Then, mount the syringe into the retention mechanism of the UD Neuroinjector and the screw to hold the plunger button.
2. Prior to first use, download the `injector control stepper` folder from the UD Neuroinjector OSE GitHub repository onto a laptop and upload the `injector control.ino` file to the Arduino UNO through the appropriate COM port using the Arduino IDE. This upload only needs to be done once. To operate the injector open a serial terminal on VS Code or the Arduino IDE at 115200 baud.
3. Select mode 4 and choose the appropriate syringe (1: Hamilton 7000.5, 0.5 µL; 2: Hamilton 7001, 1 µL; 3: custom, entered as nL mm^*−*1^); Other Hamilton or custom syringe options can be automatically written into the control script using our provided web app [17].
4. Select mode 1 for extraction and then use either the manual joystick (mode 3) or a defined extraction rate for drawing the injectable fluid into the barrel.
5. Select Mode 2 for injection and enter the desired flow rate (in nL min^*−*1^) and volume (in nL) when prompted.
6. The injection can be paused at any time using the reset push button (connected to PIN 12). The control script reports completion when the target volume has been delivered.

### 3.11 RAG and AI-driven knowledge assistance

The knowledge layer of the UD Neuroinjector OSE combines device-specific information with literature-derived stereotaxic data. Hardware and firmware information is stored and maintained in `hardware-kb.json,` while syringe specifications are stored and maintained in `hamilton-syringes.json.` These files provide information about serial communication, motor and lead-screw behavior, syringe calibration, mechanical compatibility, assembly, and troubleshooting. This is served to users using a react-based open-access web application [12].

The literature component loads several locally generated datasets, including `injection-structured.json, injection-passages.json, injection-coordinates.json, and stereotaxic-protocols.json.`The structured dataset contains LLM-extracted target coordinates, injection volumes, infusion rates, reference landmarks, and supporting source quotes. The passage dataset contains coordinate-relevant text windows extracted from methods, results, abstracts, and figure legends. The broader literature corpus provides additional paper-level context. The React client performs deterministic retrieval using lexical overlap, anatomical-region matching, PMID matching, species filtering, and coordinate-target relevance. For a coordinate query, the highest-scoring target rows are passed to the language model with their PMIDs, regions, AP/ML/DV coordinates, reference landmarks, volumes of injections, flow rates, and source quotes. Hardware and syringe records are retrieved independently and added when relevant. The language model is explicitly instructed to cite only PMIDs present in the screened corpus and to reproduce indexed values without modifications.

The application communicates with the language model through a Cloudflare Worker acting as an API proxy. The worker provides the model-completion endpoint, while the literature, hardware, syringe, and atlas data remain locally bundled with the React application.

### 3.12 Atlas integration

The assistant also includes an interactive Allen Mouse CCFv3-based atlas module implemented in `AllenAtlasViewer.jsx.` The module loads the CCFv3 annotation volume, structure ontology, coronal slices, and per-slice structure-ID maps. Users can search for a structure, select a location on a coronal slice, or enter AP, ML, and DV coordinates manually.

Atlas locations are resolved to anatomical structures using voxel-level CCFv3 structure IDs. ML coordinates are calculated relative to the atlas midline, while DV coordinates are calculated relative to the local pial surface. Atlas-derived coordinates can be sent to the assistant for comparison with literature-derived coordinates and for consistency with previously reported injection coordinates.

### 3.13 Statistical analysis

All statistical analyses were performed using custom data analysis workflows in Jupyter notebooks running Python 3.12. An ordinary least squares linear model was used for linear regression of relative meniscus displacement over time. For comparison of CV and monotonic fraction across different flow rates we used a one-way ANOVA. For comparing area of spread and DAPI^+^ cell count measurements we used two-sided Welch’s independent-samples t-test run on per-animal mean measurement.

### 3.14 Data visualization and artwork

All graphs were obtained using custom data analysis pipelines in Jupyter notebooks running python 3.12. CCFv3 coronal images were used for visualization of stereotaxic coordinates in mouse brain sections [6]. Figures were assembled using Inkscape vector graphics editor. Artwork was generated using Inkscape vector graphics editor.

## 4 Discussion

Stereotaxic neurosurgical injections are an important technique in neuroscience that requires expensive equipment, hardware, and the development of a high level of skill and precision. There is an urgent need to reduce the economic and knowledge barrier to making this technique more accessible. In this work, we have addressed the hardware, software, and knowledge barriers needed to accomplish successful stereotaxic intracranial injections in mice using the UD Neuroinjector OSE. First, the UD Neuroinjector OSE consists of a robust hardware and firmware design for an automated injector and Arduino-based control system. Using high resolution image analysis, we have shown stable and highly linear meniscus displacement rates for low volume and low flow-rate injections. Next, we have shown using *in vivo* intracranial surgery and injections into the mouse brain that the UD Neuroinjector has similar performance to a commercial injector at less than 0.1 × the cost. Finally, we have designed the UD Neuroinjector Assistant that uses AI-driven domain expertise via a RAG approach that is grounded in a literature-derived corpus. A CCFv3-based atlas helps guide and complement LLM responses and target identification assistance. All the components of the UD Neuroinjector OSE are open source. The UD Neuroinjector is the first open-source end-to-end workflow solution for intracranial injections into the mouse brain.

Automated injection of gene delivery vectors into the brain during stereotaxic surgery is a superior method compared to manual injection [18]. The approach to design automated injectors has not significantly changed since it was first systematically demonstrated in the mouse brain [18] indicating design convergence owing to the constraint of the objective–using a syringe for injection [19]. Many research groups use pressure-based approaches that use glass micropipettes for injection [1]. Multiple open-source solutions using and validating this design and approach have been reported previously [3, 20–23]. However, this approach needs the use of other expensive equipment such as pipette pullers, manual calibration of volumes, and may often lead to unused injectate due to calibration errors, which could be relatively expensive depending on the cost of the vectors. This approach also leads to reduced standardization across labs. Since syringes used for stereotaxic injections (such as Hamilton syringes) are easily sterilized, can be repeatably used for many injections, and are relatively inexpensive (∼ 100 USD–150 USD per syringe), using this setup can provide significant cost and time savings for stereotaxic injections.

Commercial automated injector systems rely on proprietary hardware design and closed-access firmware and software. Beyond the warranty period, the cost via the primary vendor or even a third-party vendor to maintain and repair these injector systems following hardware or software malfunction can be very high. Open-source integrated hardware and software systems can serve as effective solutions to this problem. Similar to our open-source design (Figure 1a–c), other recent open-source designs for automated injectors that use stepper motors have been reported [24, 25]. However, head-to-head comparison to commercial injector performance for *in vivo* injections is not available for these designs [24, 25]. Our capillary fluid displacement measurements provide a robust readout of flow-rate stability using standard operating conditions for stereotaxic intracranial injections (Fig. 2). While Bravo-Martinez et al. (2023) provide a robust design for their stereotaxic injector, they have also not provided head-to-head comparison to a commercial injector [25]. The UD Neuroinjector performs similarly to a leading commercial alternative (QSI, Stoelting Inc.) for *in vivo* performance of stereotaxic injections into the mouse brain (Fig. 3).

The open science movement has led to the formation of technology ecosystems where software, hardware, and user/developer communities interact. This approach is a welcome change from the siloed development of scientific technology and has the potential to increase sustainability. A successful example of this ecosystem is the Bonsai visual programming platform [26, 27]. The large user base of Bonsai programming has contributed to the adoption of Bonsai by commercial vendors as well [28].

Our work provides an integrated framework for the hardware and customizable software [17] needed for operation of the UD Neuroinjector. However, we also extend this framework to also provide domain expertise using a RAG-based AI-driven application (Fig. 4). Using our CCFv3-based atlas module and the RAG-based AI assistant tool, users can easily verify previously reported stereotaxic coordinates into the specific mouse brain region they are aiming to target. We have provided validation of AI-assisted target guidance for the cerebellar simplex (Fig. 4f) as a proof-of-concept. While it is likely these extracted coordinates (Fig. 4f) may not match the actual targets from the original papers on a one-to-one basis, the fact that these targets map onto the correct anatomical sub-structure within our atlas module reference system is noteworthy.

While the UD Neuroinjector possesses many advantages compared to previous open source and commercial solutions, we would like to note a few limitations. We did not opt to design a separate display and input unit for operating the UD Neuroinjector since it reduces build time and users can readily use laptops to perform this function. On the other hand, it would be beneficial to have a standalone display and input unit using low-cost hardware to allow users maximal operational flexibility. While our syringe configurator allows users to customize the control script, our hardware design prevents the use of recent syringe models. We plan to increase the number of compatible syringes in a future release. Our *in vivo* DAPI injections used three mice per injector, however, following flow-rate linearity analysis and *in vivo* validation, the UD Neuroinjector has become our standard injector-of-use for all intracranial stereotaxic injections. For the UD Neuroinjector Assistant, we plan to continue regular expansion of the stereotaxic coordinate database for the mouse brain. Additionally, the absence of rat stereotaxic coordinates represents a significant limitation. Adding a dedicated rat brain stereotaxic coordinate file and atlas module will enable users who perform rat intracranial surgeries access to the UD Neuroinjector Assistant for their workflows. Finally, while our OSE provides individual solutions to each step of the intracranial stereotaxic injection workflow, a more comprehensive and completely integrated system which can use a robotic framework to perform surgeries [29, 30] and incorporate AI and agentic capability for human-in-the-loop surgery planning would represent a significant improvement.

The UD Neuroinjector ecosystem provides a complete end-to-end solution needed to perform successful literature-validated stereotaxic intracranial injections into the mouse brain. By democratizing each step of the workflow, our work brings together a unique combination of tools that will enable users to significantly reduce the time, effort, and money required to set up this important and advanced neuroscience workflow in their lab.

## Supporting information

Online Resource 1

Online Resource 3

Manuscript LaTeX Project Zip

Online Resource 2

## Acknowledgements

We acknowledge the University of Dayton School of Engineering Multidisciplinary Design Capstone course sequence (ECE/MEE 431L-432L; Reilly Downing, Owen Beer, and Megan LaBelle were students in this course) for providing initial resources for the development of the UD Neuroinjector. We acknowledge Doug Bishop and Brian LeMaster (AI Applications and Services, University of Dayton Information Technology) for technical guidance and providing access to the University of Dayton FlyerGPT Azure resources. We also gratefully acknowledge the Ohio Supercomputer Center (OSC) for providing computational resources to Aaron Sathyanesan (Project # PNS 0489).

## Author contributions

Conceptualization: Mir Abbas Raza, Krishna Bhavithavya Kidambi, and Aaron Sathyanesan; Methodology: Mir Abbas Raza, Reilly Downing, Owen Beer, Megan LaBelle, Krishna Bhavithavya Kidambi, and Aaron Sathyanesan; Formal analysis: Mir Abbas Raza and Aaron Sathyanesan; Funding acquisition and Resources: Krishna Bhavithavya Kidambi and Aaron Sathyanesan; Investigation: Mir Abbas Raza, Sanjay Madala, and Kassidy Schroeder; Software: Reilly Downing and Aaron Sathyanesan; Project Administration and Supervision: Krishna Bhavithavya Kidambi and Aaron Sathyanesan; Visualization: Mir Abbas Raza, Sanjay Madala, Kassidy Schroeder, and Aaron Sathyanesan; Writing–original draft: Aaron Sathyanesan; Writing–review and editing: all authors.

## Declarations

The authors have no relevant financial or non-financial interests to disclose.

## Supplementary Information

*Online Resource 1* (ESM 1.csv) Spreadsheets containing ground truth values of 84 stereotaxic coordinates from Nucleus accumbens, Hippocampal CA1 region, and Caudate putamen.

*Online Resource 2* (ESM 2.mmd) Mermaid diagram of the UD Neuroinjector Assistant.

*Online Resource 3* (ESM 3.mp4) Screen recording of CCFv3-based atlas module and surgery assistant LLM operation.

## Data availability statement

Hardware design files, firmware, and web application code are publicly available in a Zenodo repository at https://doi.org/10.5281/zenodo.22871234. The experimental datasets generated and analyzed in this study will be made available to peer reviewers on request and will be deposited in a public repository upon acceptance of the manuscript.

## Notes

https://doi.org/10.5281/zenodo.22871234

### Competing Interest Statement

The authors have declared no competing interest.

## References

[1] Cetin, A., Komai, S., Eliava, M., Seeburg, P.H., Osten, P.: Stereotaxic gene delivery in the rodent brain. Nature Protocols 1, 3166–3173 (2006) 10.1038/nprot.2006.450

[2] Tye, K.M., Deisseroth, K.: Optogenetic investigation of neural circuits underlying brain disease in animal models. Nature Reviews Neuroscience 13(4), 251–266 (2012)

[3] Wijnen, B., Hunt, E.J., Anzalone, G.C., Pearce, J.M.: Open-source syringe pump library. PLOS ONE 9(9), 107216 (2014) 10.1371/journal.pone.0107216

[4] Baden, T., Chagas, A.M., Gage, G., Marzullo, T., Prieto-Godino, L.L., Euler, T.: Open labware: 3-d printing your own lab equipment. PLoS biology 13(3), 1002086 (2015)

[5] Franklin, K.B., Paxinos, G.: Paxinos and Franklin’s the Mouse Brain in Stereotaxic Coordinates, Compact: The Coronal Plates and Diagrams. Academic Press, San Diego (2019)

[6] Wang, Q., Ding, S.-L., Li, Y., Royall, J., Feng, D., Lesnar, P., Graddis, N., Naeemi, M., Facer, B., Ho, A., Dolbeare, T., Blanchard, B., Dee, N., Wakeman, W., Hirokawa, K.E., Szafer, A., Sunkin, S.M., Oh, S.W., Bernard, A., Phillips, J.W., Hawrylycz, M., Koch, C., Zeng, H., Harris, J.A., Ng, L.: The allen mouse brain common coordinate framework: A 3d reference atlas. Cell 181(4), 936–95320 (2020) 10.1016/j.cell.2020.04.007

[7] Claudi, F., Tyson, A.L., Petrucco, L., Margrie, T.W., Portugues, R., Branco, T.: Visualizing anatomically registered data with brainrender. Elife 10, 65751 (2021)

[8] Fuglstad, J.G., Saldanha, P., Paglia, J., Whitlock, J.R.: Histological e-data registration in rodent brain spaces. Elife 12, 83496 (2023)

[9] Sayers, E.: E-utilities Quick Start. In: Entrez Programming Utilities Help [Internet]. National Center for Biotechnology Information (US), Bethesda (MD) (2008). Available from: https://www.ncbi.nlm.nih.gov/books/NBK25500/. https://www.ncbi.nlm.nih.gov/books/NBK25500/

[10] Comeau, D.C., Wei, C.-H., Islamaj Doğan, R., Lu, Z.: Pmc text mining subset in bioc: about three million full-text articles and growing. Bioinformatics 35(18), 3533–3535 (2019)

[11] OpenAI: gpt-oss-120b & gpt-oss-20b Model Card (2025). https://arxiv.org/abs/2508.10925

[12] Sathyanesan, A.: UD Neuroinjector Assistant. Accessed 9 September 2026 (2026). https://asathyanesan.github.io/Neuroinjector-OSE/assistant/

[13] Raza, M.A., Downing, R., Beer, O., LaBelle, M., Madala, S., Schroeder, K., Kidambi, K.B., Sathyanesan, A.: UD Neuroinjector Open Source Ecosystem. 10.5281/zenodo.22871234. https://github.com/asathyanesan/Neuroinjector-OSE

[14] Schindelin, J., Arganda-Carreras, I., Frise, E., Kaynig, V., Longair, M., Pietzsch, T., Preibisch, S., Rueden, C., Saalfeld, S., Schmid, B., Tinevez, J.-Y., White, D.J., Hartenstein, V., Eliceiri, K., Tomancak, P., Cardona, A.: Fiji: an open-source platform for biological-image analysis. Nature Methods 9, 676–682 (2012) 10.1038/nmeth.2019

[15] Berg, S., Kutra, D., Kroeger, T., Straehle, C.N., Kausler, B.X., Haubold, C., Schiegg, M., Ales, J., Beier, T., Rudy, M., et al.: Ilastik: interactive machine learning for (bio) image analysis. Nature methods 16(12), 1226–1232 (2019)

[16] McCauley, M.: AccelStepper: AccelStepper library for Arduino. Arduino stepper motor library (2026)

[17] Sathyanesan, A.: UD Neuroinjector — Syringe Configurator. Accessed 12 September 2026 (2026). https://asathyanesan.github.io/Neuroinjector-OSE/webapp/

[18] Brooks, A.I., Halterman, M.W., Chadwick, C.A., Davidson, B.L., Haak-Frendscho, M., Radel, C., Porter, C., Federoff, H.J.: Reproducible and efficient murine CNS gene delivery using a microprocessor-controlled injector. Journal of Neuroscience Methods 80(2), 137–147 (1998) 10.1016/S0165-0270(97)00207-0

[19] Norman, D.A., Verganti, R.: Incremental and radical innovation: Design research vs. technology and meaning change. Design Issues 30(1), 78–96 (2014) 10.1162/DESIa00250

[20] Yang, Y., Atasoy, D., Su, H.H., Sternson, S.M.: Hunger states switch a flip-flop memory circuit via a synaptic AMPK-dependent positive feedback loop. Cell 146(6), 992–1003 (2011) 10.1016/j.cell.2011.07.039

[21] Booeshaghi, A.S., Beltrame, E.d.V., Bannon, D., Gehring, J., Pachter, L.: Principles of open source bioinstrumentation applied to the poseidon syringe pump system. Scientific reports 9(1), 12385 (2019)

[22] Arnold, J., Osborne, J.: Stereotaxic Injector System (2026) 10.25378/janelia.31955799.v2

[23] Dominguez, V.H., Frankfurter, M., Hayes, K.B., Kahn, M.L., Dominguez, M.H.: A portable, ultra-low cost, open-source, pedal-controlled microinjector for laboratory use. PLOS ONE 21(5), 0347487 (2026) 10.1371/journal.pone.0347487

[24] Dodson, P.: Syringe Pump. Open-source syringe pump (2018)

[25] Bravo-Martínez, J., Ortega-Tinoco, S., Garduño, J., Hernández-López, S.: Arduino based intra-cerebral microinjector device for neuroscience research. HardwareX 15, 00446 (2023) 10.1016/j.ohx.2023.e00446

[26] Lopes, G., Bonacchi, N., Frazão, J., Neto, J.P., Atallah, B.V., Soares, S., Moreira, L., Matias, S., Itskov, P.M., Correia, P.A., Medina, R.E., Calcaterra, L., Dreosti, E., Paton, J.J., Kampff, A.R.: Bonsai: an event-based framework for processing and controlling data streams. Frontiers in Neuroinformatics 9, 7 (2015) 10.3389/fninf.2015.00007

[27] Lopes, G., Monteiro, P.: New open-source tools: Using bonsai for behavioral tracking and closed-loop experiments. Frontiers in Behavioral Neuroscience 15, 647640 (2021) 10.3389/fnbeh.2021.647640

[28] MBF Biosciences: NeuroPhotoMetrics FP3002. Accessed 16 September 2026 (2026). https://web.archive.org/web/20260628175230/ https://www.mbfbioscience.com/products/fp3002

[29] Rynes, M.L., Ghanbari, L., Schulman, D.S., Linn, S., Laroque, M., Dominguez, J., Navabi, Z.S., Sherman, P., Kodandaramaiah, S.B.: Assembly and operation of an open-source, computer numerical controlled (cnc) robot for performing cranial microsurgical procedures. Nature protocols 15(6), 1992–2023 (2020)

[30] Coffey, K.R., Barker, D.J., Ma, S., West, M.O.: Building an open-source robotic stereotaxic instrument. Journal of Visualized Experiments (80), 51006 (2013) 10.3791/51006

