## Supplementary material for "An integrated stereotaxic injection architecture for mouse intracranial surgeries with AI-driven domain expertise": Manuscript LaTeX Project Zip: Raza_et_al_Neuroinjector.pdf

$$\texttt{Displacement Resolution} = \frac{0.6096\,\mathrm{mm}}{1600\,\mathrm{microsteps}} = 0.381\,\mu\mathrm{m/microstep} \qquad (1)$$

An Arduino UNO board controls the NEMA 8 motor and TMC2208 stepper motor driver (Fig. 1c). An analog joystick controls manual injector up/down movement. A limit switch (Fig. 1c) checks if the plunger has been fully depressed and, if switched

**Table 1 Accuracy of LLM-distilled coordinates by brain region and by stereotaxic axis.**

| Accuracy by brain region | | | Accuracy by stereotaxic axis | | |
|---|---|---|---|---|---|
| **Region** | **Score** | **Percentage** | **Axis** | **Score** | **Percentage** |
| Nucleus accumbens (Acb) | 23/23 | 100% | Antero-posterior (AP) | 83/84 | 98.8% |
| Hippocampal CA1 region (CA1) | 27/29 | 93.1% | Medio-lateral (ML) | 84/84 | 100% |
| Caudate putamen (CP) | 32/32 | 100% | Dorso-ventral (DV) | 83/84 | 98.8% |

For analysis, we used features on frame-by-frame line intensity scans that covered both the black reference band as well as the meniscus (similar to Fig. 2a–b). We first sought to reduce the effect of camera jitter by calculating the vertical axis coordinate of the meniscus, $y_{\text{meniscus}}(t)$, relative to the black band reference on the capillary, $y_{\text{band}}(t)$, for $N \approx 200$ sampled data points per injection:

$$y_{\text{rel}}(t) = y_{\text{meniscus}}(t) - y_{\text{band}}(t) \tag{2}$$

We fitted displacement over time data using an ordinary least squares linear model. We then calculated the residual coefficient of variation $CV$:

$$CV = \frac{RMSE}{\overline{y}_{\text{rel}}} = \frac{\sqrt{\frac{1}{N}\sum_{i=1}^{N}\Big(y_{\text{rel}}(t_i) - \hat{y}_{\text{rel}}(t_i)\Big)^2}}{\frac{1}{N}\sum_{i=1}^{N} y_{\text{rel}}(t_i)} \tag{3}$$

Where $\hat{y}_{\text{rel}}(t_i)$ is linear regression fit of $y_{\text{rel}}$ at time $t_i$. We also calculated monotonic fraction, $f_{\text{mono}}$, as:

$$f_{\text{mono}} = \frac{1}{N-1}\sum_{i=1}^{N-1} I\left(\Delta y_{\text{smooth},i} \geq -\delta\right) \tag{4}$$

Where:

- $y_{\text{smooth}}(t_i) = \text{median}\left(y_{\text{rel}}(t_{i-3}), \ldots, y_{\text{rel}}(t_{i+3})\right)$
- $\Delta y_{\text{smooth},i} = y_{\text{smooth}}(t_{i+1}) - y_{\text{smooth}}(t_i)$
- $\delta = 0.02 \times \text{median}\left(|\Delta y_{\text{smooth},i}|\right)$
- $I(\cdot) = 1$ if true, 0 otherwise.

### 3.7 UD Neuroinjector 3-D printing and assembly

The UD Neuroinjector assembly needs low-cost off-the-shelf components and 3D-printed parts. We used a standard FFF (or FDM) printer to 3D-print the parts for the UD Neuroinjector. We have provided detailed documentation for step-by-step assembly of the UD Neuroinjector on the UD Neuroinjector OSE GitHub Repository [13]. The time needed to build the UD Neuroinjector once all the parts have been assembled is $\sim$ 30 minutes. We have also included all source STL files for 3D-printed parts as well as a bill of materials.

The default motor housing design helps secure the UD Neuroinjector onto the stereotaxic arm on the left-hand side of the stereotaxic frame. However, in the STL folder of the GitHub repository we have also provided a file (`Motor Housing_righthanded.STL`) that can be used to 3D-print a motor housing that can be secured to a “right” stereotaxic arm if needed.

### 3.8 Arduino-based actuation

We used an Arduino UNO R3 microcontroller board with a prototype expansion shield for controlling the UD Neuroinjector. A NEMA 8 bipolar motor (1.8° step angle, 28 mm body length, 0.2 A rated current, 1.6 N cm holding torque; Amazon, Seattle, WA, United States) coupled to an ultra-precision $\frac{1}{8}$-inch lead screw (41.7 threads $\text{in}^{-1}$, 0.6096 mm pitch; McMaster-Carr, Elmhurst, IL, United States) via an aluminum rigid shaft coupler ($\frac{1}{8}$-inch-to-5 mm; Amazon, Seattle, WA, United States) provided linear actuation.

To increase stability and smoothness of operation, the plunger depresser is mounted onto an MGN12H carriage block that slides along a 250 mm linear guide rail (Amazon, Seattle, WA, United States). The full steps per revolution, $N_{\text{step}}$, of the NEMA 8 are calculated as:

$$N_{\text{step}} = \frac{360°}{1.8°/\text{step}} = 200 \text{ steps/revolution.} \tag{5}$$

A TMC2208 stepper motor driver (Amazon, Seattle, WA, United States) enables microstepping. Microsteps per revolution, $N_{\text{micro}}$, and degrees per microstep, $\theta_{\text{micro}}$, are calculated as:

$$N_{\text{micro}} = 200 \textit{ full steps} \times 8 \textit{ microsteps per step} = 1600 \textit{ microsteps per turn} \tag{6}$$

$$\theta_{\text{micro}} = \frac{1.8}{8} = 0.225° \textit{ per microstep} \tag{7}$$

Linear actuation per turn, $Actuation_{\text{turn}}$, i.e. for 360°, considering lead screw dimensions ($\frac{1}{8}$-inch) is 0.6096 mm. Linear actuation for step ($Actuation_{\text{step}}$), and microstep ($Actuation_{\text{micro}}$), is calculated as:

$$\text{Actuation}_{\text{step}} = \frac{\text{Actuation}_{\text{turn}}}{200} = \frac{0.6096}{200} = 0.003\,048\,\text{mm}\ (3.048\,\mu\text{m}) \tag{8}$$

$$\text{Actuation}_{\text{micro}} = \frac{\text{Actuation}_{\text{turn}}}{\frac{200}{1/8}} = \frac{0.6096}{1600} = 0.000\,381\,\text{mm}\ (0.381\,\mu\text{m}) \tag{9}$$

#### 3.9 Firmware and serial programming

The Arduino control script (`injector_control.ino`) is built on the open-source AccelStepper library [16], driving the TMC2208 through a step/dir interface (digital pins 8 and 9) over a 115200-baud serial link. The firmware implements four operating modes: extraction, injection, manual joystick control, and syringe selection, chosen by serial commands. Motion is executed as relative displacement from a reference position that is re-zeroed at the start of each mode. Syringe selection configures the plunger gradation $g$, defined by the firmware variable `syringe_div`, in nanoliters of fluid displaced per millimeter of plunger travel. Compile-time constants define two presets: `HAM_7000_5_DIV` $= 25/3 \approx 8.33\,\text{nL}\,\text{mm}^{-1}$ for the Hamilton 7000.5 (0.5 µL; PN: 86250, Hamilton Company, Reno, NV, United States) syringe and `HAM_7001_DIV` $= 50/3 \approx 16.67\,\text{nL}\,\text{mm}^{-1}$ for the Hamilton 7001 (1 µL; PN: 80100, Hamilton Company, Reno, NV, United States) syringe. Selecting a syringe (mode 4) copies the corresponding constant into `syringe_div`; a custom gradation ($1\,\text{nL}\,\text{mm}^{-1}$ to $1000\,\text{nL}\,\text{mm}^{-1}$) can be entered for any other syringe. The presets correspond to the full syringe volume delivered over the 60 mm plunger travel of the Hamilton 700-series microliter syringes (e.g., $8.33\,\text{nL}\,\text{mm}^{-1} \times 60\,\text{mm} = 500\,\text{nL}$ for the 0.5 µL syringe).

For a commanded injection or extraction volume $V$ (nL), the required plunger displacement is converted to microsteps as:

$$\text{steps} = \frac{V \times N_{\text{micro}}}{g \times L} \tag{10}$$

where $L$ is the lead screw pitch (firmware variable `lead`, 0.6096 mm revolution$^{-1}$) and $N_{\text{micro}}$ = 1600 microsteps revolution$^{-1}$ (firmware constant `stepsprev`). For a requested flow rate $F$ (nL min$^{-1}$), the firmware computes the required microstep rate, $S$, as:

$$S = \frac{F \times N_{\text{micro}}}{60 \times g \times L} \tag{11}$$

In the firmware code, Equations 10 and 11 are evaluated as `steps = vol * stepsprev / (syringe_div * lead)` and `steppsec = (flow / syringe_div) / lead / 60 * stepsprev`, respectively. The firmware validates all user input before motion begins, accepting flow rates of 0.1 nL min$^{-1}$ to 1000 nL min$^{-1}$ and volumes of 0.1 nL to 500 nL; the upper volume bound corresponds to the full stroke of the 0.5 µL syringe. For example, a 100 nL injection at 100 nL min$^{-1}$ using the 0.5 µL syringe corresponds to 31,496 microsteps (12.0 mm of plunger travel) executed at 524.9 microsteps s$^{-1}$, completed in 60 s. The table below (Table 2) maps the notation to its firmware representation.

**Table 2 Mapping of mathematical notation to firmware representation.**

| Symbol | Firmware representation | Meaning |
|---|---|---|
| $g$ | `syringe_div` (set from `HAM_7000_5_DIV`, `HAM_7001_DIV`, or custom entry) | Plunger gradation, nL mm$^{-1}$ |
| $N_{\text{micro}}$ | `stepsprev` | microsteps per revolution (1600) |
| $L$ | `lead` | lead screw pitch, 0.6096 mm revolution$^{-1}$ |
| $V$ | `vol` | commanded volume, nL |
| $F$ | `flow` | commanded flow rate, nL min$^{-1}$ |
| $S$ | `steppsec` | commanded microstep rate, microsteps s$^{-1}$ |
| steps | `step` | commanded displacement, microsteps |

[19] Norman, D.A., Verganti, R.: Incremental and radical innovation: Design research vs. technology and meaning change. Design Issues **30**(1), 78–96 (2014) https://doi.org/10.1162/DESI_a_00250

[20] Yang, Y., Atasoy, D., Su, H.H., Sternson, S.M.: Hunger states switch a flip-flop memory circuit via a synaptic AMPK-dependent positive feedback loop. Cell **146**(6), 992–1003 (2011) https://doi.org/10.1016/j.cell.2011.07.039

[21] Booeshaghi, A.S., Beltrame, E.d.V., Bannon, D., Gehring, J., Pachter, L.: Principles of open source bioinstrumentation applied to the poseidon syringe pump system. Scientific reports **9**(1), 12385 (2019)

[22] Arnold, J., Osborne, J.: Stereotaxic Injector System (2026) https://doi.org/10.25378/janelia.31955799.v2

[23] Dominguez, V.H., Frankfurter, M., Hayes, K.B., Kahn, M.L., Dominguez, M.H.: A portable, ultra-low cost, open-source, pedal-controlled microinjector for laboratory use. PLOS ONE **21**(5), 0347487 (2026) https://doi.org/10.1371/journal.pone.0347487

[24] Dodson, P.: Syringe Pump. Open-source syringe pump (2018)

[25] Bravo-Martínez, J., Ortega-Tinoco, S., Garduño, J., Hernández-López, S.:

Arduino based intra-cerebral microinjector device for neuroscience research. HardwareX **15**, 00446 (2023) https://doi.org/10.1016/j.ohx.2023.e00446

[26] Lopes, G., Bonacchi, N., Frazão, J., Neto, J.P., Atallah, B.V., Soares, S., Moreira, L., Matias, S., Itskov, P.M., Correia, P.A., Medina, R.E., Calcaterra, L., Dreosti, E., Paton, J.J., Kampff, A.R.: Bonsai: an event-based framework for processing and controlling data streams. Frontiers in Neuroinformatics **9**, 7 (2015) https://doi.org/10.3389/fninf.2015.00007

[27] Lopes, G., Monteiro, P.: New open-source tools: Using bonsai for behavioral tracking and closed-loop experiments. Frontiers in Behavioral Neuroscience **15**, 647640 (2021) https://doi.org/10.3389/fnbeh.2021.647640

[28] MBF Biosciences: NeuroPhotoMetrics FP3002. Accessed 16 September 2026 (2026). https://web.archive.org/web/20260628175230/https://www.mbfbioscience.com/products/fp3002

[29] Rynes, M.L., Ghanbari, L., Schulman, D.S., Linn, S., Laroque, M., Dominguez, J., Navabi, Z.S., Sherman, P., Kodandaramaiah, S.B.: Assembly and operation of an open-source, computer numerical controlled (cnc) robot for performing cranial microsurgical procedures. Nature protocols **15**(6), 1992–2023 (2020)

[30] Coffey, K.R., Barker, D.J., Ma, S., West, M.O.: Building an open-source robotic stereotaxic instrument. Journal of Visualized Experiments (80), 51006 (2013) https://doi.org/10.3791/51006

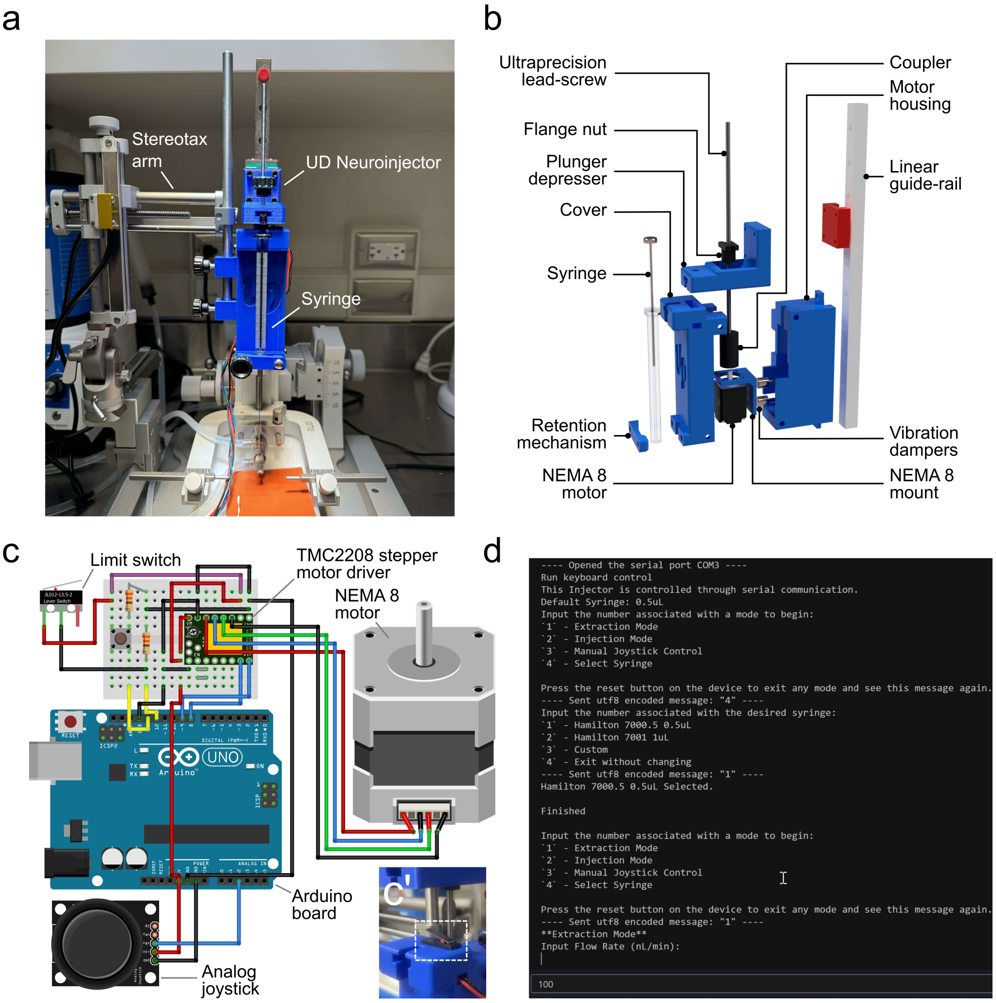


**Fig. 1 UD Neuroinjector Hardware Design and Control. a.** A photograph of the UD Neuroinjector mounted onto the arm of a mouse stereotaxic setup. **b.** Exploded view of the components of the UD Neuroinjector **c.** Wiring diagram (fritzing) of Arduino microcontroller-based control of the NEMA 8 motor in the UD Neuroinjector; inset (**c'**) shows limit switch placement to prevent damage. **d.** Serial-programming-based interface for controlling the UD Neuroinjector. The script for UD Neuroinjector control and serial programming can be downloaded from the UD Neuroinjector GitHub repository.

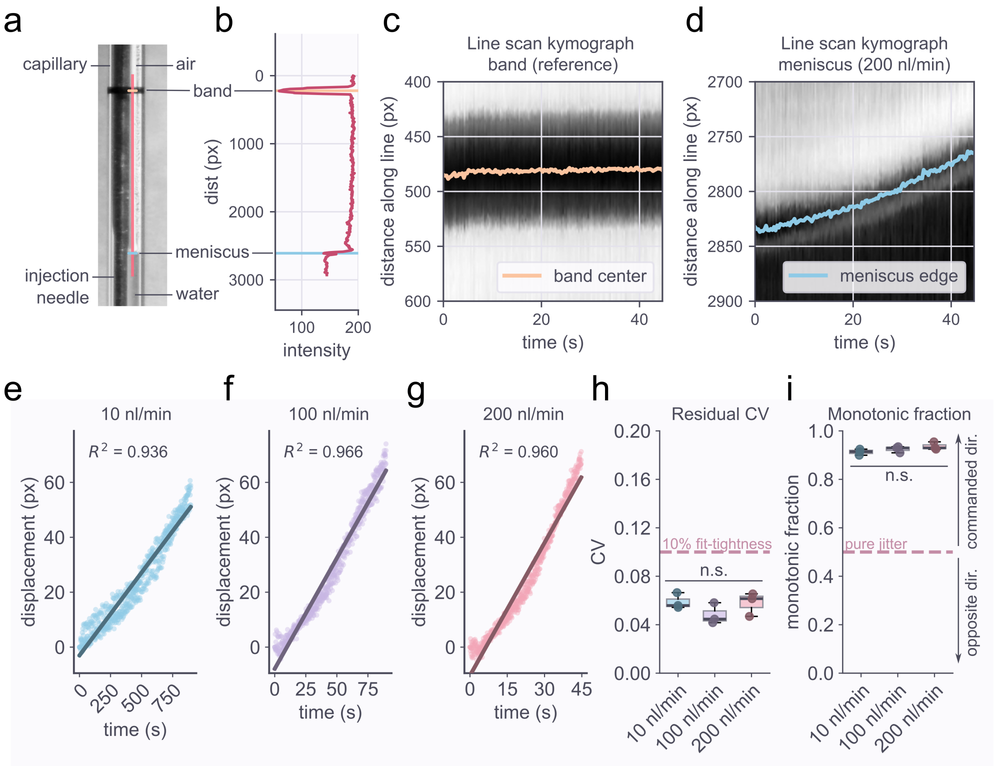


**Fig. 2 Flow rate validation for UD Neuroinjector. a.** Fluid capillary displacement setup showing the black band ('band') reference, syringe injection needle, and the water meniscus at the air-water interface inside the capillary. **b.** Line intensity scan profile of the orange line in panel a showing the inverted peak (orange) corresponding to the reference black band as well as the meniscus profile peak (blue). **c.** Line scan kymographs for the center of the black reference band (orange) and **d.** the meniscus edge (blue) for a representative 200 nl/min injection. **e.** Meniscus relative displacement over the course of the injection for 10 nl/min **f.** 100 nl/min and **g.** 200 nl/min flow rate. **h.** Residual CV comparison between the different flow rate measurements (no statistically significant difference [n.s.] between CV across flow rates; N = 3 injections per group; One-way ANOVA, P = 0.3043). **i.** Comparison of monotonic fraction across different flow rates (no statistically significant difference [n.s.] between monotonic fraction across flow rates; N = 3 injections per group; One-way ANOVA, P = 0.2071)

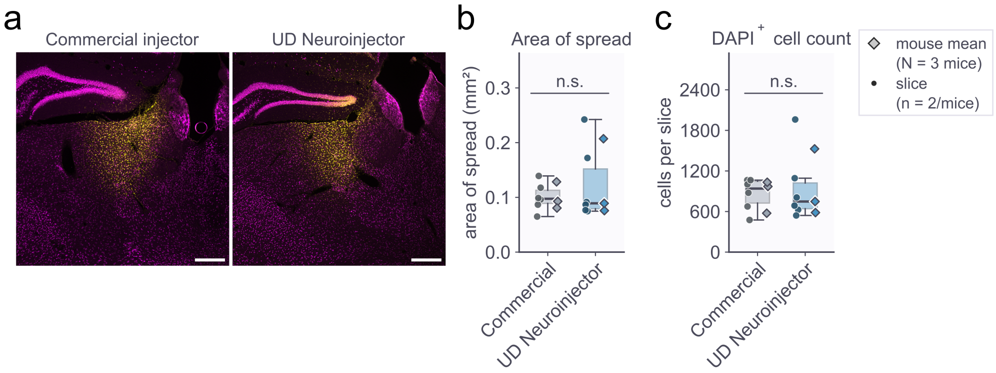


**Fig. 3** ***In vivo*** **head-to-head performance of UD Neuroinjector vs. a commercial injector.** **a.** Representative fluorescence microscopy images of mouse brain tissue sections following stereotaxic surgery and injection of 100 nl of 5 mg/ml DAPI solution using the Quintessential Stereotaxic Injector (QSI, Stoelting Company) (left image) and the UD Neuroinjector (right image) (AP $\sim -1.2$ to $-1.4$ mm, ML $\sim 0.9$ to 1.2 mm, DV $\sim -2.6$ to $-2.8$ mm); DAPI shown in yellow, anti-NeuN pan-neuronal marker shown in magenta. **b.** Area of spread comparison between the commercial injector and the UD Neuroinjector shows no statistically significant (n.s.) differences in area of spread ($N = 3$ mice, $n = 2$ slices/mouse; Welch's independent-samples t-test on per-animal mean measurement; Welch's $t = -0.5313$, two-sided $P = 0.6392$). **c.** No statistically significant differences in DAPI+ cell counts ($N = 3$ mice, $n = 2$ slices/mouse; Welch's independent-samples t-test; Welch's $t = -0.2889$, two-sided $P = 0.792$) between the commercial injector and the UD Neuroinjector. Scale bar for panel a = 150 µm.

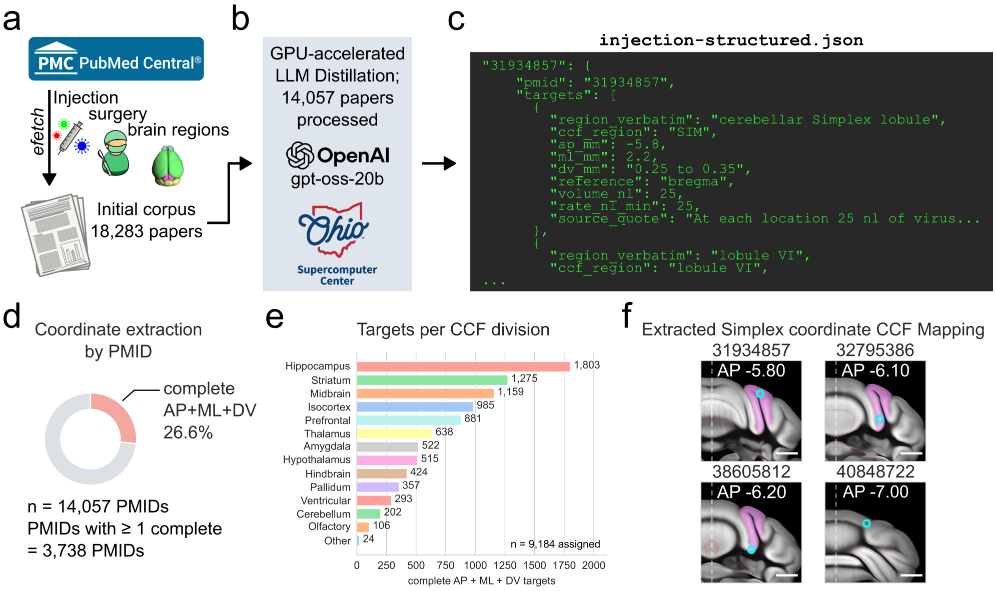


**Fig. 4 Corpus-grounded AI-driven domain expertise and knowledge layer for the UD Neuroinjector ecosystem. a.** Initial scoping of PubMed Central (PMC) for search terms associated with injection, surgery, and mouse brain regions yielded an initial corpus of 18,283 papers. **b.** We used GPU-accelerated distillation of the papers using the `gpt-oss-20b` open source model on the Ohio Supercomputer and obtained **c.** a structured file, `injection-structured.json`. **d.** The `injection-structured.json` file contains 3,738 PMIDs with at least one complete (antero-posterior – AP, mediolateral – ML, and dorsoventral – DV) coordinate set representing 26.6% of the 14,057 PMIDs. **e.** An analysis of the targets per common coordinate framework (CCF) division (n = 9,184 targets assigned) reveals the spread of the coordinates, with the hippocampus having the largest number of previously reported coordinates followed by the striatum. **f.** Proof-of-feasibility mapping of extracted coordinates (cyan circles) onto the CCFv3 coordinate system for the cerebellar simplex (magenta) found in PMIDs 31934857 (AP -5.80 relative to bregma), 32795386 (AP -6.10 relative to bregma), 38605812 (AP -6.20; relative to bregma but rederived from lambda – 4.20), and 40848722 (AP -7.00; relative to bregma but rederived from lambda – 4.20).
