## Supplementary material for "An integrated stereotaxic injection architecture for mouse intracranial surgeries with AI-driven domain expertise": Manuscript LaTeX Project Zip: Raza_et_al_Neuroinjector.tex

%Version 3.1 December 2024
% See section 11 of the User Manual for version history
%
%%%%%%%%%%%%%%%%%%%%%%%%%%%%%%%%%%%%%%%%%%%%%%%%%%%%%%%%%%%%%%%%%%%%%%
%%                                                                 %%
%% Please do not use \input{...} to include other tex files.       %%
%% Submit your LaTeX manuscript as one .tex document.              %%
%%                                                                 %%
%% All additional figures and files should be attached             %%
%% separately and not embedded in the \TeX\ document itself.       %%
%%                                                                 %%
%%%%%%%%%%%%%%%%%%%%%%%%%%%%%%%%%%%%%%%%%%%%%%%%%%%%%%%%%%%%%%%%%%%%%

%%\documentclass[referee,sn-basic]{sn-jnl}% referee option is meant for double line spacing

%%=======================================================%%
%% to print line numbers in the margin use lineno option %%
%%=======================================================%%

%%\documentclass[lineno,pdflatex,sn-basic]{sn-jnl}% Basic Springer Nature Reference Style/Chemistry Reference Style

%%=========================================================================================%%
%% the documentclass is set to pdflatex as default. You can delete it if not appropriate.  %%
%%=========================================================================================%%

%%\documentclass[sn-basic]{sn-jnl}% Basic Springer Nature Reference Style/Chemistry Reference Style

%%Note: the following reference styles support Namedate and Numbered referencing. By default the style follows the most common style. To switch between the options you can add or remove Numbered in the optional parenthesis. 
%%The option is available for: sn-basic.bst, sn-chicago.bst%  
 
%%\documentclass[pdflatex,sn-nature]{sn-jnl}% Style for submissions to Nature Portfolio journals
%%\documentclass[pdflatex,sn-basic]{sn-jnl}% Basic Springer Nature Reference Style/Chemistry Reference Style
\documentclass[pdflatex,sn-mathphys-num]{sn-jnl}% Math and Physical Sciences Numbered Reference Style
%%\documentclass[pdflatex,sn-mathphys-ay]{sn-jnl}% Math and Physical Sciences Author Year Reference Style
%%\documentclass[pdflatex,sn-aps]{sn-jnl}% American Physical Society (APS) Reference Style
%%\documentclass[pdflatex,sn-vancouver-num]{sn-jnl}% Vancouver Numbered Reference Style
%%\documentclass[pdflatex,sn-vancouver-ay]{sn-jnl}% Vancouver Author Year Reference Style
%%\documentclass[pdflatex,sn-apa]{sn-jnl}% APA Reference Style
%%\documentclass[pdflatex,sn-chicago]{sn-jnl}% Chicago-based Humanities Reference Style

%%%% Standard Packages
%%<additional latex packages if required can be included here>

\usepackage{graphicx}%
\graphicspath{{figures/}}
\usepackage{adjustbox}
\usepackage{multirow}%
\usepackage{amsmath,amssymb,amsfonts}%
\usepackage{amsthm}%
\usepackage{mathrsfs}%
\usepackage[title]{appendix}%
\usepackage{xcolor}%
\usepackage{textcomp}%
\usepackage{manyfoot}%
\usepackage{booktabs}%
\usepackage{algorithm}%
\usepackage{algorithmicx}%
\usepackage{algpseudocode}%
\usepackage{listings}%
\usepackage{tabularray}
\usepackage{siunitx} % This is the units package
\DeclareSIUnit{\rev}{rev} 
\DeclareSIUnit{\revolution}{revolution}
\DeclareSIUnit{\microstep}{microstep}
\DeclareSIUnit{\microsteps}{microsteps}
\DeclareSIUnit{\frame}{frame}
\DeclareSIUnit{\frames}{frames}
\DeclareSIUnit{\threads}{threads}
\DeclareSIUnit{\inch}{in}
\DeclareSIUnit{\steps}{steps} 
%%%%
%\usepackage[backend=biber,style=ieee,sorting=none,giveninits=true,maxbibnames=99,doi=false,
%url=false,eprint=false,isbn=false]{biblatex}
%
%\addbibresource{sn-bibliography}
%% as per the requirement new theorem styles can be included as shown below
\theoremstyle{thmstyleone}%
\newtheorem{theorem}{Theorem}%  meant for continuous numbers
%%\newtheorem{theorem}{Theorem}[section]% meant for sectionwise numbers
%% optional argument [theorem] produces theorem numbering sequence instead of independent numbers for Proposition
\newtheorem{proposition}[theorem]{Proposition}% 
%%\newtheorem{proposition}{Proposition}% to get separate numbers for theorem and proposition etc.

\theoremstyle{thmstyletwo}%
\newtheorem{example}{Example}%
\newtheorem{remark}{Remark}%

\theoremstyle{thmstylethree}%
\newtheorem{definition}{Definition}%

\raggedbottom
%%\unnumbered% uncomment this for unnumbered level heads

\newcounter{onlineresource}
\newcommand{\olr}[1]{%
  \refstepcounter{onlineresource}%
  \label{#1}%
  Online Resource \theonlineresource%
}

\begin{document}

\title[An integrated stereotaxic injection architecture for mouse intracranial surgeries with AI-driven domain expertise
]{An integrated stereotaxic injection architecture for mouse intracranial surgeries with AI-driven domain expertise}
	

%%=============================================================%%
%% GivenName	-> \fnm{Joergen W.}
%% Particle	-> \spfx{van der} -> surname prefix
%% FamilyName	-> \sur{Ploeg}
%% Suffix	-> \sfx{IV}
%% \author*[1,2]{\fnm{Joergen W.} \spfx{van der} \sur{Ploeg} 
%%=============================================================%%
 
\author[1]{\fnm{} \sur{Mir Abbas Raza}} 
\author[2]{\fnm{Reilly} \sur{Downing}} 
\author[3]{\fnm{Owen} \sur{Beer}} 
\author[3]{\fnm{Megan} \sur{LaBelle}} 
\author[1]{\fnm{Sanjay} \sur{Madala}} 
\author[1]{\fnm{Kassidy} \sur{Schroeder}} 
\author[3]{\fnm{Krishna Bhavithavya} \sur{Kidambi}} 
}

%\equalcont{These authors contributed equally to this work.}

\affil[1]{\orgdiv{Department of Biology, College of Arts \& Sciences}, \orgname{University of Dayton}, \orgaddress{\street{300 College Park}, \city{Dayton}, \postcode{45469}, \state{OH}, \country{USA}}}

\affil[2]{\orgdiv{Department of Electrical \& Computer Engineering, School of Engineering}, \orgname{University of Dayton}, \orgaddress{\street{300 College Park}, \city{Dayton}, \postcode{45469}, \state{OH}, \country{USA}}}

\affil[3]{\orgdiv{Department of Mechanical and Aerospace Engineering, School of Engineering}, \orgname{University of Dayton}, \orgaddress{\street{300 College Park}, \city{Dayton}, \postcode{45469}, \state{OH}, \country{USA}}}

%%==================================%%
%% Sample for unstructured abstract %%
%%==================================%%

\abstract{Stereotaxic intracranial microinjection in mice is a critical step in neuroscience workflows, enabled by robust hardware design, precise operation, and neuroanatomical expertise. Manual injections are prone to error and stereotax-mountable commercial automated injectors are extremely expensive. Further, the knowledge needed for performing accurate intracranial injections and surgeries is heterogeneous and disparate. Here we demonstrate the design and characterization of the UD Neuroinjector software-hardware-knowledge architecture, which serves to both reduce the cost of performing precisely controlled stereotaxic microinjections and provide a web platform for planning accurate targeting of specific mouse brain regions. We designed the UD Neuroinjector using low-cost hardware and 3D-printed parts, with a total build-cost of $\sim$\$300 --- less than $\text{0.1}\times$ the cost of a commercial injector. Arduino-based firmware enables programmable flow rates, automated injection and extraction, and manual joystick control. Capillary-fluid measurements show stable, highly linear displacement ($\text{R}^2>\text{0.93}$) at commonly used injection rates. In a head-to-head comparison with a commercial injector, \textit{in vivo} DAPI injections into the mouse thalamus showed comparable spread and $\text{DAPI}^+$ cell counts. For effective experimental design of intracranial injections, the web-based components of our architecture provide AI-enabled assistance and atlas-guided navigation. We used a hybrid retrieval-augmented generation (RAG) approach to ground AI-assisted protocol guidance for mouse intracranial surgeries using a corpus of 3,738 papers. Using the Common Coordinate Framework version 3 (CCFv3), we designed an interactive tool for identifying target coordinates. Together, the UD Neuroinjector architecture integrates and enables efficient knowledge discovery, experimental design, planning, and implementation of mouse intracranial surgeries.}
 

\keywords{stereotaxic, injector, intracranial surgery, mouse, brain atlas}

%%\pacs[JEL Classification]{D8, H51}

%%\pacs[MSC Classification]{35A01, 65L10, 65L12, 65L20, 65L70}

\maketitle

\section{Introduction}\label{sec1}
Stereotaxic microinjections into the rodent brain are a fundamental part of a wide range of neuroscience experiments. The efficiency of approaches such as neuroanatomical tracing, induced cell ablation studies, and viral induction of biosensors for \textit{in vivo} physiology or optogenetics largely depend on stable injection of chemicals or biologics at precise flow rates into the brain \cite{Cetin2006Stereotaxic,tye2012optogenetic}. While manual stereotaxic injections into the rodent brain are still commonly practiced, neuroscience research labs often resort to using commercial automated microinjection systems. These commercial injector systems are very expensive (upwards of $\$$3000) and can be cost-prohibitive in resource-poor contexts \cite{Wijnen2014SyringePump,baden2015open}. However, few open-access and low-cost solutions for automated stereotaxic microinjectors exist. Importantly, head-to-head \textit{in vivo} validation in comparison to commercial injectors is lacking.

In addition to the automation of intracranial injections, researchers performing these injections also rely on knowledge components such as mouse or rat brain atlases and previously published protocols \cite{franklin2019paxinos,Wang2020Allen}. Given the sheer anatomical and cellular complexity combined with the small size of the mouse brain, the precision of injection into a particular brain region depends on the accurate identification of stereotaxic coordinates of the said region. While few recent tools have been designed for planning viral injections, these are most appropriate for planning neural recording probe trajectories which require additional considerations of pitch and yaw angles in 3D space \cite{claudi2021visualizing,fuglstad2023histological}. For stereotaxic viral injections in general, the first main consideration includes the identification of target coordinates along the antero-posterior (AP), mediolateral (ML), and dorsoventral (DV) axes using a standardized coronal brain atlas. The second main consideration includes prior knowledge of successful stereotaxic targeting from peer-reviewed publications. However, no tool has incorporated the rich corpus of stereotaxic neurosurgical information from previously published studies in the mouse.

We designed an end-to-end open-source architecture -- the UD Neuroinjector Open Source Ecosystem (OSE) -- that includes hardware, software, and knowledge components that can be applied across the entire workflow of stereotaxic intracranial injection into the mouse brain. We built the UD Neuroinjector using fused filament fabrication 3D printed parts, low-cost electronic and mechanical components, and simple serial control via an Arduino microcontroller board. The UD Neuroinjector is capable of stable flow rates that are commonly used in mouse intracranial injection protocols. \textit{In vivo} intracranial DAPI injections showed that the UD Neuroinjector is similar in performance to that of a commercial injector. Finally, we used a hybrid retrieval augmented generation (RAG) approach to build domain expertise in mouse brain surgery. This information is easily accessible through an LLM via a chat-based interface. Using the Allen Brain Atlas Common Coordinate Framework version 3 (CCFv3), we designed a simple web app to complement domain expertise through AI-assisted protocol planning. Our system enables comprehensive workflow planning, troubleshooting, and implementation of successful stereotaxic intracranial injections into the mouse brain, which is a cornerstone technique in modern neuroscience.
%%%%%%%%%%%%%%%%%%%%%%%%%%%%%%%%%%%%%%%%%%%%%%%%%%%%%%%%%%%%%%%%%%%
 
\section{Results}\label{sec2}

\subsection{Hardware and software layer}

\subsubsection{UD Neuroinjector design and control}
We used off-the-shelf electronic and mechanical components combined with 3D-printed mounting and housing for the UD Neuroinjector, with a net cost
of 289 USD (Fig.~\ref{fig1}a-b). We used a NEMA 8 bipolar stepper motor (1.8$^\circ$ full step angle; 200 full steps per revolution, i.e., 360$^\circ$) coupled to an ultraprecision $\frac{1}{8}$ inch lead screw (0.6096 \unit{\mm\per\rev}) to drive a flange nut connected to a 3D-printed plunger depresser that slides along a linear guide rail (Fig.~\ref{fig1}b).

A 3D-printed NEMA 8 mount holds the motor in place within the motor housing with vibration dampers for stable operation. The outer part of the cover for the motor housing has a recess to hold the upper lip of
the syringe barrel in place, with the lower part of the barrel secured to the cover using a retention mechanism
(Fig.~\ref{fig1}b). This helps hold the syringe securely in place during injections. The plunger depresser recess is designed to hold the plunger button in place so it can move precisely along the vertical axis. The side of the motor housing facing the stereotaxic arm has two locations along which the standard-size stereotaxic arm adapter can be passed through to mount the UD Neuroinjector (Fig.~\ref{fig1}a). A pair of set screws secure the injector to the stereotaxic arm adaptor.

The NEMA 8 stepper motor is driven by a TMC2208 stepper motor driver (Fig.~\ref{fig1}c), which is configured for
microstepping yielding 1600 microsteps per revolution. Given this mechanical configuration, the theoretical displacement resolution is calculated as follows:

\begin{equation}
	\texttt{Displacement Resolution} = \frac{0.6096 \: \si{\mm}}{1600 \: \si{\microsteps}}  = 0.381 \: \si{\mu m \per\microstep}
		\label{eq:displacement_resolution}
\end{equation} 

An Arduino UNO board controls the NEMA 8 motor and TMC2208 stepper motor
driver (Fig.~\ref{fig1}c). An analog joystick
controls manual injector up/down movement. A limit switch
(Fig.~\ref{fig1}c) checks if the plunger has been
fully depressed and, if switched ON, prevents further stepping. A
simple, interactive, serial programming interface controls the injector
(Fig.~\ref{fig1}d).


\subsubsection{Flow rate validation for UD Neuroinjector}

Since intracranial injections into the mouse brain depend on stable and relatively low flow rates, we sought to determine how stably the UD Neuroinjector can dispense a defined volume (150~nl) at different flow
rates commonly used in mouse intracranial injections. Using a high-resolution macro video camera, we recorded water meniscus displacement in a 50~$\mu$l capillary tube to measure injection flow rate stability (Fig.~\ref{fig2}a). Since water is injected into the capillary, the meniscus is displaced upward towards a black band (reference) marked on the capillary tube.

Using automated line intensity scan analysis, we tracked the movement
of the meniscus (Fig.~\ref{fig2}a--b, blue line) with respect
to the reference band (Fig.~\ref{fig2}a--b, orange line).
Representative line scan kymographs show no noticeable drift in the
position of the black reference band (Fig.~\ref{fig2}c) and a
steady upward movement of the meniscus edge
(Fig.~\ref{fig2}d).

The UD Neuroinjector showed stable linear meniscus displacement at injection
rates of 10~nl/min (Fig.~\ref{fig2}e; $R^2=0.936$),
\qty{100}{\nano\liter\per\minute} (Fig.~\ref{fig2}f; $R^2=0.966$), and 200~nl/min
(Fig.~\ref{fig2}g; $R^2=0.960$). Fit tightness using residual
CV did not show significant statistical differences across different
flow rates (Fig.~\ref{fig2}h).

We also measured monotonic fraction as a measure of fractional forward
progression over a rolling window. Mean monotonic fraction exceeded
0.9 for all injection flow rates tested (Fig.~\ref{fig2}i).
We also did not notice any statistically significant difference in
monotonic fraction across flow rates, indicating stable forward
injection across conditions. For every injection, the plunger moved
completely to the bottom of the barrel, indicating that the entire
150~nl had been dispensed.


\subsubsection{\textit{In vivo} head-to-head performance of the UD Neuroinjector}

To perform head-to-head comparison of the UD Neuroinjector with a
commercial stereotax-mountable injector (Quintessential Stereotaxic
Injector, Stoelting Co., Wood Dale, IL, United States), we injected the
DNA-intercalating fluorescent stain
4$'$,6-diamidino-2-phenylindole (DAPI) into the mouse lateral posterior thalamic nucleus (AP $\sim -1.2$~ to $-1.4$ mm, ML $\sim 0.9$~ to $1.2$ mm, DV $\sim -2.6$~ to $-2.8$ mm) just ventral
to the medial dentate gyrus (Fig.~\ref{fig3}a).

Computational image analysis of DAPI staining showed that the UD
Neuroinjector performed comparably to the commercial injector both in
terms of the total area of spread across the injected region
(Fig.~\ref{fig3}b; Welch's t-test on per-mouse measurements; no statistically
significant difference between groups; $N=3$ mice per group; Welch's t = -0.53; df = 2.47; P = 0.64) as well as the automated DAPI$^+$
cell counts (Fig.~\ref{fig3}c; Welch's t-test on per-mouse measurements; no statistically
significant difference between groups; $N=3$ mice per group; Welch's t = -0.29; df = 2.92; P = 0.79). The UD Neuroinjector
thus has similar performance as a standard automated commercial injector in
terms of \textit{in vivo} intracranial injections.

\subsubsection{Routine laboratory use}
In everyday practice, we have used the UD Neuroinjector to inject adeno-associated viral (AAV) vectors into the thalamus and cerebellum at multiple stereotaxic coordinates. A retrospective analysis of histologically confirmed tissue data in our lab between September 1, 2025, and May 12, 2026, showed that the UD Neuroinjector had a success rate of 8/10 injections, similar to the commercial injector with a success rate of 9/10 injections that showed intended expression in the target brain region. In cases of the failed injections for both injectors we could not accurately determine the cause of failure in terms of histological expression.

\subsection{Domain-expertise and knowledge layer}

Stereotaxic injections into the mouse brain are challenging at both
the hardware-software level as well as the knowledge and domain-expertise
level. Having provided a stable and robust open-source solution using
the UD Neuroinjector described above, we proceeded to design the
knowledge and domain-expertise component necessary for successful
intracranial injections into the mouse brain
(Fig.~\ref{fig4}).


\subsubsection{Initial corpus scoping}

We developed a Node.js/JavaScript trawler pipeline
(\texttt{ingest-protocols.js}) that queried the NCBI PubMed and PMC
databases using the NCBI E-utilities \texttt{efetch} endpoint \cite{sayers2008quickstart} for mouse
stereotaxic injections into brain regions. This broad-based approach
ensured that we would get maximal coverage of the available corpus,
retrieving article metadata and available full text. When
\texttt{efetch} was unable to extract full text, we used the PMC BioC
API \cite{comeau2019pmc} as a fallback.

We focused on passages that included the results, text from the figure
captions and legends, and the methods section. Following deduplication,
this text was assembled into the \texttt{surgical-protocols.json} file.
This JSON file has stereotaxic-surgery-related text for a total of
18,283 papers (Fig.~\ref{fig4}a).


\subsubsection{LLM-based corpus distillation}

To sort through these large passages of text and identify mouse brain
stereotaxic coordinates, we used an LLM-based approach on the Ohio
Supercomputer (Fig.~\ref{fig4}b). First, we used
keyword-based filtering to identify papers with mouse, stereotaxic,
coordinate, and injection-related signals and build vLLM batch prompt
files. We then used a distillation SLURM script on OSC to run OpenAI's
open-weight model \texttt{gpt-oss-20b} (21B parameters)
\cite{OpenAI2025GPTOSS} over the prompts.

The final outputs were collected into the
\texttt{injection-structured.json} file
(Fig.~\ref{fig4}c), which contains coordinates (at least
one type---AP, ML, or DV---extracted) for 14,057 unique PMIDs, of which
26.6\% (3,738 PMIDs) have at least one complete set (AP, ML, and DV)
(Fig.~\ref{fig4}d). A total of 13,044 target regions
were extracted, from which 9,184 were confidently assigned to a
particular CCF division (Fig.~\ref{fig4}e), with the
hippocampus and striatum comprising a large percentage of the targets. We independently curated a ground-truth dataset (\olr{res:csv}) for targets previously reported in the Nucleus Accumbens (Acb), Hippocampal CA1 region (CA1), and the Caudate putamen (CP). We used this dataset to test the accuracy of the LLM-extracted coordinates (Table~\ref{tab:accuracy}). In comparison to ground-truth, the average accuracy of LLM-distilled stereotaxic coordinates by brain region was 97.7\% (Acb: 100\%; CA1: 93.1\%; CP: 100\%). The average accuracy by stereotaxic coordinate axis was 99.2\% (AP: 98.8\%; ML: 100\%; DV: 98.8\%).

\begin{table}[htbp]
\caption{\textbf{Accuracy of LLM-distilled coordinates by brain region and by stereotaxic axis.}}
\label{tab:accuracy}
\centering
\footnotesize
\begin{tabular}{
    @{}
    >{\centering\arraybackslash}p{0.20\textwidth}
    >{\centering\arraybackslash}p{0.09\textwidth}
    >{\centering\arraybackslash}p{0.11\textwidth}
    @{\hspace{2em}}
    >{\centering\arraybackslash}p{0.20\textwidth}
    >{\centering\arraybackslash}p{0.09\textwidth}
    >{\centering\arraybackslash}p{0.11\textwidth}
    @{}
}
\toprule
\multicolumn{3}{c}{\textbf{Accuracy by brain region}}
& \multicolumn{3}{c}{\textbf{Accuracy by stereotaxic axis}} \\
\cmidrule(r){1-3} \cmidrule(l){4-6}
\textbf{Region} & \textbf{Score} & \textbf{Percentage}
& \textbf{Axis} & \textbf{Score} & \textbf{Percentage} \\
\midrule
Nucleus accumbens (Acb)
& 23/23 & 100\%
& Antero-posterior (AP)
& 83/84 & 98.8\% \\
Hippocampal CA1 region (CA1)
& 27/29 & 93.1\%
& Medio-lateral (ML)
& 84/84 & 100\% \\
Caudate putamen (CP)
& 32/32 & 100\%
& Dorso-ventral (DV)
& 83/84 & 98.8\% \\
\botrule
\end{tabular}
\end{table}


\subsubsection{Identification of mouse brain stereotaxic coordinates using RAG}

We implemented a simple UD Neuroinjector Assistant chat module that
uses deterministic retrieval to identify stereotaxic coordinates
grounded in the scientific corpus (\olr{res:mmd}) \cite{Sathyanesan2026Assistant,Raza_UD_Neuroinjector_Open_2026}. The UD
Neuroinjector Assistant uses a React client to load the structured
coordinates and relevant extracted passages. The system tokenizes user
input and ranks candidate targets and literature records based on
lexical similarity, anatomical-region matching, PMID match, and
coordinate-target relevance. Regional queries are also given a
``structure-aware'' score.

To ensure maximal fidelity to mouse-specific stereotaxic coordinates,
a mouse-specific filter was enforced. Highest-scoring candidates
receive priority in the output response, which contains the PMID,
anatomical region as stated or CCFv3-matched, AP/ML/DV coordinates,
injection volume, flow rate, and the original source quote. Strict
grounding rules are enforced for the language model (GPT 5.5 or GPT
5.4) to only cite papers found in
\texttt{injection-structured.json}.

Additionally, to close the hardware-software-knowledge loop, we also
made hardware troubleshooting and syringe information relevant to the
UD Neuroinjector available to the assistant. The UD Neuroinjector
Assistant thus serves as a ``one-stop-shop'' for stereotaxic target
planning using a RAG-based approach with deterministic lexical and
anatomical-context matching.


\subsubsection{A simple CCFv3-based atlas module for target coordinate planning}

We designed a simple and interactive atlas module using the Allen Mouse
Common Coordinate Framework version 3 (CCFv3) for effective stereotaxic
target planning \cite{Sathyanesan2026Assistant}. The module uses the
CCFv3 and the Allen structure ontology to map voxels onto anatomical
structures and region names. We use coronal slices indexed by AP axis
position and overlaid with anatomical boundaries. Users can search for
a particular location using a dropdown or click-to-target a particular
brain region.

For a selected location on a brain region within a defined AP slice,
ML and DV coordinates are calculated and displayed in millimeters
relative to the CCFv3 reference axes. Targets can also be resolved
manually by users. Once a target with AP, ML, and DV (w.r.t. Bregma) coordinates has
been identified, this can be queried by the LLM to check for
consistency with previously published papers, as seen in the screen-recording (\olr{res:mp4}).

Using the atlas in combination with corpus-grounded AI-driven domain
expertise may serve as an effective strategy for optimal stereotaxic
targeting. We verified the accuracy of extracted coordinates for the
cerebellar simplex (or simple lobule) from specific papers (unique
PMIDs) by mapping them onto the closest-matching AP slice images in our
atlas module reference system (Fig.~\ref{fig4}f). We
observed striking concordance for the extracted coordinates
(Fig.~\ref{fig4}f, cyan circle) within the simplex
(Fig.~\ref{fig4}f, magenta shaded region).

%%%%%%%%%%%%%%%%%%%%%%%%%%%%%%%%%%%%%%%%%%%%%%%%%%%%%%%%%%%%%%%%%%%%%
\section{Materials and Methods}\label{sec:methods}

\subsection{Animals}\label{subsec:animals}

All animal procedures were performed according to the Institutional
Animal Care and Use Committee of the University of Dayton
(Protocol 023-04) and the Guide for the Care and Use of Laboratory
Animals (NIH). We used adult C57BL/6 and B6EiC3Sn.BLiAF1/J mice in
injection experiments. Both male and female sexes were used for
injection experiments.

\subsection{Surgery and stereotaxic intracranial injections}
\label{subsec:surgery}

Mice were weighed and subjected to anesthesia induction
(3\% Isoflurane) with isoflurane in a chamber linked to a table-top
isoflurane system (RWD Life Science Inc., Dover, DE, United States).
After induction, mice were placed on a temperature-controlled heating
stage (ALA Scientific, Farmingdale, NY, United States). Body temperature
was monitored using a rectal probe connected to the heating stage to
maintain optimal body temperature. The snout was then positioned inside
an anesthesia mask (RWD Life Science Inc., Dover, DE, United States)
and 3\% Isoflurane was supplied through this line.

The mouse was then positioned carefully in the digital stereotaxic frame
(RWD Life Science Inc., Dover, DE, United States) by placing the ear bars
slowly into the ear canal. Lubricant ointment (Paralube vet ointment,
Dechra, Boston, MA, United States) was topically applied to the eyes to
prevent drying. Once the breathing rate stabilized (1--1.5 breaths per
second), isoflurane was reduced to $\sim1.5\%$ -- 2.5\% for anesthesia maintenance.

The fur above the skull was closely clipped using surgical scissors.
Betadine was applied to the skin above the skull using a sterile cotton
swab. 0.5\% Lidocaine (2.5--4 mg/kg; Vedco, St. Joseph, MO, United
States) was injected locally in the skin above the skull as a local
anesthetic. A single incision was made along the midline to expose the
skull using a disposable scalpel (Foster City, CA, United States).
Bleeding if any was stemmed using Sugi\textsuperscript{\textregistered}
Sponge Points (Kettenbach, Huntington Beach, CA, United States).
Sterile cotton swabs were used to clear away any tissue above the skull.

Once the skull was clearly visible, 3\% hydrogen peroxide was applied
to the skull to visualize the bregma. Saline was then immediately
applied to neutralize the hydrogen peroxide. Excess solution was cleared
away using sterile cotton swabs. The bregma was then recorded digitally,
following which a 0.6 mm burr hole was drilled in the skull at
AP $\sim -1.2$~ to $-1.4$ mm, ML $\sim 0.9$~ to $1.2$ mm,  using an electric dental drill.
Sterile saline was applied to the holes to prevent drying. The dura was carefully removed using fine forceps. 100 nl of 5 mg/ml DAPI solution (10.9 mM solution reconstituted in saline; BioLegend, San Diego, CA, United States) was injected into the
thalamus at a depth (DV) of $\sim -2.6$~ to $-2.8$ mm using a 1 $\mu$l syringe
(7001KH, PN: 80100, Hamilton Company, Reno, NV, United States) mounted on an
automated injector (either the UD Neuroinjector or the Quintessential
Stereotaxic Injector, Stoelting Co., Wood Dale, IL, United States) at a
flow rate of 100 nl/min.

Two minutes after the DAPI solution had been completely injected into the
site, the syringe was gently retracted. Vetbond tissue glue
(3M, St. Paul, MN, United States) was applied to glue the skin flaps
back. Meloxicam (5 mg/kg) was injected into the scruff for analgesia,
and isoflurane was gradually reduced. The mouse was allowed to rest
on the stage till it regained consciousness and generated active
movement. It was then transferred to its home cage and monitored for
1 hour following completion of surgery. The mouse was monitored once every 12 hours for signs of pain, dehydration, reduction of activity and discomfort. The mouse was then transcardially perfused within 24 hours after surgery, and the brain was used for tissue processing and immunohistochemistry. We performed one injection per mouse into one hemisphere.


\subsection{Tissue processing and immunohistochemistry}
\label{subsec:tissue-processing}

Mice injected with DAPI were anesthetized with isoflurane, and following
tailpinch-verification, transcardially perfused with ice-cold 0.1 M
Phosphate Buffered Saline (PBS) followed by 4\%
paraformaldehyde (PFA). The brain was dissected out of the skull and
post-fixed in 4\% PFA with 30\% Sucrose overnight at \qty{4}{\degreeCelsius} after which the
brain was stored in 0.1 M PBS with 30\% sucrose at \qty{4}{\degreeCelsius} till the tissue
processing stage.

The brain was cut into free-floating 35 $\mu$m coronal sections using
a freezing-stage (BFS-40MPA, Physitemp Instruments, Clifton, NJ, United
States) mounted on a sliding microtome (HM 430, Epredia, Kalamazoo, MI,
United States). Sections were labeled using standard
immunohistochemical procedures.

To visualize neurons in DAPI-stained sections, we used an anti-NeuN
primary antibody (1:2000; Cat. No. 266 004, Synaptic Systems GmbH,
Goettingen, Germany) followed by an Alexa Fluor 488 anti-guinea-pig
secondary antibody (Code: 706-545-148, Jackson Immunoresearch Laboratories,
West Grove, PA, United States). The tissue was then mounted onto Fisherbrand
Superfrost Plus Microscope slides (Fisher Scientific Company, Hanover
Park, IL, United States), with a few drops of Fluoromount-G
(SouthernBiotech, Birmingham, AL, United States) and covered with a
cover glass (Fisher Scientific Company, Hanover Park, IL, United States).


\subsection{Fluorescence microscopy and imaging}
\label{subsec:fluorescence-imaging}

We used an Olympus BX53 epifluorescence microscope equipped with an LED
illumination system, DAPI/FITC/Cy3/Cy5 filters, an Olympus DP23M
monochrome camera, and Olympus CellSens software (Evident Scientific
Inc., Needham, MA, United States) for fluorescence imaging of DAPI$^+$
and NeuN$^+$ neurons. Images were captured using a 4$\times$ Olympus
air objective in the TIFF format using the CellSens software at
3088$\times$2076 pixel resolution.


\subsection{Automated cell quantification}
\label{subsec:cell-quantification}

We used ImageJ Fiji \cite{Schindelin2012Fiji} and the pixel
classification tool Ilastik \cite{Berg2019Ilastik} for automated
DAPI$^+$ cell quantification. We converted the TIFF microscopy images
to PNG and used these files as input training data to train an Ilastik
segmentation model. Ilastik simple segmentation files were then analyzed
on Fiji using simple thresholding. For DAPI$^+$ cell counting we used
the `Analyze particles' tool. For calculating area of spread we first
applied a gaussian blur followed by thresholding to binarize the area with DAPI$^+$ cells and the rest of the brain section area with no DAPI. After this we used the ‘Analyze particles’ tool to quantify the area of spread in mm$^2$.

\subsection{Fluid displacement video acquisition and analysis}
\label{subsec:fluid-displacement}

We inserted a Hamilton syringe injection needle (Hamilton 7000.5,
PN: 86250, Hamilton Company, Reno, NV, United States) into a
\qty{50}{\micro\litre} glass capillary tube (Yankee Disposable Micropet,
BD, Parsippany, NJ, United States) pre-filled with distilled water such
that there is little air space between the black reference band on the
capillary and the water meniscus. The bottom of the capillary was sealed
off with parafilm. The capillary was held upright using a clamp. We used
a Samsung Galaxy S25 Ultra (Sony IMX854 camera sensor) mounted in a secure
holder for acquiring ultrahigh definition video at
\qty{30}{\frames\per\second} and $25\times$ magnification for a pixel
resolution of $3840\times2160$.

For analysis, we used features on frame-by-frame line intensity scans
that covered both the black reference band as well as the meniscus
(similar to Fig.~\ref{fig2}a--b). We first sought to reduce the effect of camera
jitter by calculating the vertical axis coordinate of the meniscus,
$y_{\mathrm{meniscus}}(t)$, relative to the black band reference on the
capillary, $y_{\mathrm{band}}(t)$, for $N \approx 200$ sampled data points
per injection:

\begin{equation}
y_{\mathrm{rel}}(t)
=
y_{\mathrm{meniscus}}(t)
-
y_{\mathrm{band}}(t)
\label{eq:relative-meniscus}
\end{equation}

We fitted displacement over time data using an ordinary least squares
linear model. We then calculated the residual coefficient of variation
$CV$:

\begin{equation}
CV
=
\frac{RMSE}{\overline{y}_{\mathrm{rel}}}
=
\frac{
\sqrt{
\displaystyle
\frac{1}{N}
\sum_{i=1}^{N}
\bigg(
y_{\mathrm{rel}}(t_i)
-
\hat{y}_{\mathrm{rel}}(t_i)
\bigg)^2
}
}{
\displaystyle
\frac{1}{N}
\sum_{i=1}^{N}
y_{\mathrm{rel}}(t_i)
}
\label{eq:residual-cv}
\end{equation}

Where $\hat{y}_{\mathrm{rel}}(t_i)$ is linear regression fit of
$y_{\mathrm{rel}}$ at time $t_i$. We also calculated monotonic fraction,
$f_{\mathrm{mono}}$, as:

\begin{equation}
f_{\mathrm{mono}}
=
\frac{1}{N-1}
\sum_{i=1}^{N-1}
I\left(
\Delta y_{\mathrm{smooth},i} \geq -\delta
\right)
\label{eq:monotonic-fraction}
\end{equation}

Where:
\begin{itemize}
    \item
    $y_{\mathrm{smooth}}(t_i)
    =
    \operatorname{median}
    \left(
    y_{\mathrm{rel}}(t_{i-3}),\ldots,
    y_{\mathrm{rel}}(t_{i+3})
    \right)$

    \item
    $\Delta y_{\mathrm{smooth},i}
    =
    y_{\mathrm{smooth}}(t_{i+1})
    -
    y_{\mathrm{smooth}}(t_i)$

    \item
    $\delta
    =
    0.02 \times
    \operatorname{median}
    \left(
    \left|\Delta y_{\mathrm{smooth},i}\right|
    \right)$

    \item
    $I(\cdot)=1$ if true, 0 otherwise.
\end{itemize}

\subsection{UD Neuroinjector 3-D printing and assembly}
\label{subsec:printing-assembly}

The UD Neuroinjector assembly needs low-cost off-the-shelf components
and 3D-printed parts. We used a standard FFF (or FDM) printer to 3D-print
the parts for the UD Neuroinjector. We have provided detailed
documentation for step-by-step assembly of the UD Neuroinjector on the
UD Neuroinjector OSE GitHub Repository
\cite{Raza_UD_Neuroinjector_Open_2026}. The time needed to build the UD Neuroinjector once all the parts have been assembled is $\sim30$ minutes. We have also included all source STL files for
3D-printed parts as well as a bill of materials.

The default motor housing design helps secure the UD Neuroinjector onto
the stereotaxic arm on the left-hand side of the stereotaxic frame.
However, in the STL folder of the GitHub repository we have also
provided a file (\texttt{Motor Housing\_righthanded.STL}) that can be
used to 3D-print a motor housing that can be secured to a ``right''
stereotaxic arm if needed. 

\subsection{Arduino-based actuation}
\label{subsec:arduino-actuation}

We used an Arduino UNO R3 microcontroller board with a prototype
expansion shield for controlling the UD Neuroinjector. A NEMA 8 bipolar
motor (\ang{1.8} step angle, \qty{28}{\milli\metre} body length,
\qty{0.2}{\ampere} rated current, \qty{1.6}{\newton\centi\metre}
holding torque; Amazon, Seattle, WA, United States) coupled to an
ultra-precision $\frac{1}{8}$-inch lead screw
(\qty{41.7}{\threads\per\inch}, \qty{0.6096}{\milli\metre} pitch;
McMaster-Carr, Elmhurst, IL, United States) via an aluminum rigid
shaft coupler ($\frac{1}{8}$-inch-to-\qty{5}{\milli\metre};
Amazon, Seattle, WA, United States) provided linear actuation.

To increase stability and smoothness of operation, the plunger
depresser is mounted onto an MGN12H carriage block that slides along
a \qty{250}{\milli\metre} linear guide rail (Amazon, Seattle, WA,
United States). The full steps per revolution, $N_{\mathrm{step}}$,
of the NEMA 8 are calculated as:

\begin{equation}
N_{\mathrm{step}}
=
\frac{\ang{360}}{\ang{1.8}/\mathrm{step}}
=
200 \:\: \si{\steps/\revolution}.
\label{eq:full-steps}
\end{equation}

A TMC2208 stepper motor driver (Amazon, Seattle, WA, United States)
enables microstepping. Microsteps per revolution, $N_{\mathrm{micro}}$,
and degrees per microstep, $\theta_{\mathrm{micro}}$, are calculated as:

\begin{equation}
N_{\mathrm{micro}}
=
200~\textit{full steps}
\times
8~\textit{microsteps per step}
=
1600~\textit{microsteps per turn}
\label{eq:microsteps-per-revolution}
\end{equation}

\begin{equation}
\theta_{\mathrm{micro}}
=
\frac{1.8}{8}
=
\ang{0.225}~\textit{per microstep}
\label{eq:degrees-per-microstep}
\end{equation}

Linear actuation per turn, $Actuation_{\mathrm{turn}}$, i.e.\
for \ang{360}, considering lead screw dimensions
($\frac{1}{8}$-inch) is \qty{0.6096}{\milli\metre}.
Linear actuation for step ($Actuation_{\mathrm{step}}$), and
microstep ($Actuation_{\mathrm{micro}}$), is calculated as:

\begin{equation}
\mathrm{Actuation_{step}}
=
\mathrm{\frac{Actuation_{turn}}{200}}
=
\frac{0.6096}{200}
=
\qty{0.003048}{\milli\metre}
\;(\qty{3.048}{\micro\metre})
\label{eq:actuation-step}
\end{equation}

\begin{equation}
\mathrm{Actuation_{micro}}
=
\mathrm{\frac{Actuation_{turn}}
{\frac{200}{1/8}}}
=
\frac{0.6096}{1600}
=
\qty{0.000381}{\milli\metre}
\;(\qty{0.381}{\micro\metre})
\label{eq:actuation-micro}
\end{equation}

\subsection{Firmware and serial programming}
\label{subsec:firmware-serial}

The Arduino control script (\texttt{injector\_control.ino}) is built on the
open-source AccelStepper library \cite{McCauley2026AccelStepper}, driving the
TMC2208 through a step/dir interface (digital pins 8 and 9) over a
115200-baud serial link. The firmware implements four operating modes:
extraction, injection, manual joystick control, and syringe selection,
chosen by serial commands. Motion is executed as relative displacement from
a reference position that is re-zeroed at the start of each mode. Syringe
selection configures the plunger gradation $g$, defined by the firmware
variable \texttt{syringe\_div}, in nanoliters of fluid displaced per
millimeter of plunger travel. Compile-time constants define two presets:
\texttt{HAM\_7000\_5\_DIV} = $25/3 \approx
\qty{8.33}{\nano\litre\per\milli\metre}$ for the Hamilton 7000.5
(\qty{0.5}{\micro\litre}; PN: 86250, Hamilton Company, Reno, NV,
United States) syringe and \texttt{HAM\_7001\_DIV} = $50/3 \approx
\qty{16.67}{\nano\litre\per\milli\metre}$ for the Hamilton 7001
(\qty{1}{\micro\litre}; PN: 80100, Hamilton Company, Reno, NV,
United States) syringe. Selecting a syringe (mode 4) copies the
corresponding constant into \texttt{syringe\_div}; a custom gradation
(\qtyrange{1}{1000}{\nano\litre\per\milli\metre}) can be entered for
any other syringe. The presets correspond to the full syringe volume
delivered over the \qty{60}{\milli\metre} plunger travel of the Hamilton
700-series microliter syringes (e.g.,
$\qty{8.33}{\nano\litre\per\milli\metre}
\times \qty{60}{\milli\metre}
= \qty{500}{\nano\litre}$ for the \qty{0.5}{\micro\litre} syringe).

For a commanded injection or extraction volume $V$ (\unit{\nano\litre}),
the required plunger displacement is converted to microsteps as:

\begin{equation}
\mathrm{steps}
=
\frac{V \times N_{\mathrm{micro}}}{g \times L}
\label{eq:volume-to-microsteps}
\end{equation}

where $L$ is the lead screw pitch (firmware variable \texttt{lead},
\qty{0.6096}{\milli\metre\per\revolution}) and
$N_{\mathrm{micro}} = \qty{1600}{\microsteps\per\revolution}$
(firmware constant \texttt{stepsprev}). For a requested flow rate $F$
(\unit{\nano\litre\per\minute}), the firmware computes the required
microstep rate, $S$, as:

\begin{equation}
S
=
\frac{F \times N_{\mathrm{micro}}}
{60 \times g \times L}
\label{eq:microstep-rate}
\end{equation}

In the firmware code, Equations~\ref{eq:volume-to-microsteps} and
\ref{eq:microstep-rate} are evaluated as
\texttt{steps = vol * stepsprev / (syringe\_div * lead)} and
\texttt{steppsec = (flow / syringe\_div) / lead / 60 * stepsprev},
respectively. The firmware validates all user input before motion begins,
accepting flow rates of
\qtyrange{0.1}{1000}{\nano\litre\per\minute} and volumes of
\qtyrange{0.1}{500}{\nano\litre}; the upper volume bound corresponds
to the full stroke of the \qty{0.5}{\micro\litre} syringe. For example,
a \qty{100}{\nano\litre} injection at
\qty{100}{\nano\litre\per\minute} using the
\qty{0.5}{\micro\litre} syringe corresponds to 31,496 microsteps
(\qty{12.0}{\milli\metre} of plunger travel) executed at
\qty{524.9}{\microsteps\per\second}, completed in
\qty{60}{\second}. The table below (Table~\ref{tab:firmware_notation}) maps the notation to its firmware representation.

\begin{table}[htbp]
\caption{\textbf{Mapping of mathematical notation to firmware representation.}}
\label{tab:firmware_notation}
\centering
\footnotesize

\begin{tabular}{
	@{}
	>{\centering\arraybackslash}p{0.10\textwidth}
	>{\raggedright\arraybackslash}p{0.4\textwidth}
	>{\raggedright\arraybackslash}p{0.4\textwidth}
	@{}
}
\toprule
\textbf{Symbol}
& \textbf{Firmware representation}
& \textbf{Meaning} \\
\midrule

$g$
& \texttt{syringe\_div} (set from \texttt{HAM\_7000\_5\_DIV},
  \texttt{HAM\_7001\_DIV}, or custom entry)
& Plunger gradation, \unit{\nano\litre\per\milli\metre} \\

$N_{\mathrm{micro}}$
& \texttt{stepsprev}
& microsteps per revolution (1600) \\

$L$
& \texttt{lead}
& lead screw pitch, \qty{0.6096}{\milli\metre\per\revolution} \\

$V$
& \texttt{vol}
& commanded volume, \unit{\nano\litre} \\

$F$
& \texttt{flow}
& commanded flow rate, \unit{\nano\litre\per\minute} \\

$S$
& \texttt{steppsec}
& commanded microstep rate, \unit{\microsteps\per\second} \\

$\mathrm{steps}$
& \texttt{step}
& commanded displacement, microsteps \\

\botrule
\end{tabular}
\end{table}
 
\subsection{Typical operation}
\label{subsec:typical-operation}

Plunger displacement per microstep determines volumetric resolution of the
UD Neuroinjector:

\begin{equation}
\Delta V_{\mathrm{micro}}
=
\frac{g \times L}{N_{\mathrm{micro}}}
\label{eq:volumetric-resolution}
\end{equation}

Where $\Delta V_{\mathrm{micro}}$ is the volume delivered per microstep,
which for a Hamilton 7000.5 (\qty{0.5}{\micro\litre}) syringe is
$\approx \qty{0.0032}{\nano\litre}$ (\qty{3.2}{\pico\litre}) per
microstep and $\approx \qty{0.0064}{\nano\litre}$
(\qty{6.4}{\pico\litre}) for the Hamilton 7001
(\qty{1}{\micro\litre}) syringe. The achievable flow-rate range is
determined by the maximum step rate
(\qty{8000}{\microsteps\per\second}). The accepted input range of
\qtyrange{0.1}{1000}{\nano\litre\per\minute} maps to
\qtyrange{0.53}{5249}{\microsteps\per\second} on the
\qty{0.5}{\micro\litre} syringe
(\qtyrange{0.26}{2625}{\microsteps\per\second} on the
\qty{1}{\micro\litre} syringe). The control script also reports
percentage injection progress at \qty{1}{\hertz}.

In the manual joystick mode, Analog PIN A2 translates up-or-down (X)
displacement from the joystick into a
\qty{600}{\nano\litre\per\minute} plunger stroke in the corresponding
direction for fast control. The downward plunger stroke is gated by the
limit switch (PIN 12).

Follow the instructions below for a standard injection:

\begin{enumerate}
    \item Mount the
    injector onto the stereotaxic arm. Then, mount the syringe into the retention mechanism of the UD Neuroinjector and the screw to hold the plunger button.

    \item Prior to first use, download the \texttt{injector\_control\_stepper} folder from the UD Neuroinjector OSE GitHub repository onto a laptop and upload the \texttt{injector\_control.ino} file to the Arduino UNO through the appropriate COM port using the Arduino IDE. This upload only needs to be done once. To operate the injector open a serial terminal on VS Code or the Arduino IDE at 115200 baud.

    \item Select mode 4 and choose the appropriate syringe
    (1: Hamilton 7000.5, \qty{0.5}{\micro\litre};
    2: Hamilton 7001, \qty{1}{\micro\litre};
    3: custom, entered as \unit{\nano\litre\per\milli\metre});
    Other Hamilton or custom syringe options can be automatically
    written into the control script using our provided web app
    \cite{Sathyanesan2026Configurator}.

    \item Select mode 1 for extraction and then use either the manual
    joystick (mode 3) or a defined extraction rate for drawing the
    injectable fluid into the barrel.

    \item Select Mode 2 for injection and enter the desired flow rate
    (in \unit{\nano\litre\per\minute}) and volume
    (in \unit{\nano\litre}) when prompted.

    \item The injection can be paused at any time using the reset push
    button (connected to PIN 12). The control script reports completion
    when the target volume has been delivered.
\end{enumerate}

\subsection{RAG and AI-driven knowledge assistance}
\label{subsec:rag-ai}

The knowledge layer of the UD Neuroinjector OSE combines device-specific
information with literature-derived stereotaxic data. Hardware and firmware
information is stored and maintained in \texttt{hardware-kb.json}, while
syringe specifications are stored and maintained in
\texttt{hamilton-syringes.json}. These files provide information about
serial communication, motor and lead-screw behavior, syringe calibration,
mechanical compatibility, assembly, and troubleshooting. This is served to
users using a react-based open-access web application
\cite{Sathyanesan2026Assistant}.

The literature component loads several locally generated datasets, including
\texttt{injection-structured.json}, \texttt{injection-passages.json},
\texttt{injection-coordinates.json}, and
\texttt{stereotaxic-protocols.json}. The structured dataset contains
LLM-extracted target coordinates, injection volumes, infusion rates,
reference landmarks, and supporting source quotes. The passage dataset
contains coordinate-relevant text windows extracted from methods, results,
abstracts, and figure legends. The broader literature corpus provides
additional paper-level context.

The React client performs deterministic retrieval using lexical overlap,
anatomical-region matching, PMID matching, species filtering, and
coordinate-target relevance. For a coordinate query, the highest-scoring
target rows are passed to the language model with their PMIDs, regions,
AP/ML/DV coordinates, reference landmarks, volumes of injections, flow
rates, and source quotes. Hardware and syringe records are retrieved
independently and added when relevant. The language model is explicitly
instructed to cite only PMIDs present in the screened corpus and to
reproduce indexed values without modifications.

The application communicates with the language model through a Cloudflare
Worker acting as an API proxy. The worker provides the model-completion
endpoint, while the literature, hardware, syringe, and atlas data remain
locally bundled with the React application.


\subsection{Atlas integration}
\label{subsec:atlas-integration}

The assistant also includes an interactive Allen Mouse CCFv3-based atlas
module implemented in \texttt{AllenAtlasViewer.jsx}. The module loads the
CCFv3 annotation volume, structure ontology, coronal slices, and per-slice
structure-ID maps. Users can search for a structure, select a location on
a coronal slice, or enter AP, ML, and DV coordinates manually.

Atlas locations are resolved to anatomical structures using voxel-level
CCFv3 structure IDs. ML coordinates are calculated relative to the atlas
midline, while DV coordinates are calculated relative to the local pial
surface. Atlas-derived coordinates can be sent to the assistant for
comparison with literature-derived coordinates and for consistency with
previously reported injection coordinates.

\subsection{Statistical analysis}
\label{subsec:statistical-analysis}

All statistical analyses were performed using custom data analysis
workflows in Jupyter notebooks running Python 3.12. An ordinary least
squares linear model was used for linear regression of relative meniscus
displacement over time. For comparison of CV and monotonic fraction across
different flow rates we used a one-way ANOVA. For comparing area of spread
and DAPI$^+$ cell count measurements we used two-sided Welch's
independent-samples t-test run on per-animal mean measurement.


\subsection{Data visualization and artwork}
\label{subsec:data-visualization}

All graphs were obtained using custom data analysis pipelines in Jupyter
notebooks running python 3.12. CCFv3 coronal images were used for
visualization of stereotaxic coordinates in mouse brain sections
\cite{Wang2020Allen}. Figures were assembled using Inkscape vector
graphics editor. Artwork was generated using Inkscape vector graphics
editor.


\section{Discussion}
\label{sec:discussion}

Stereotaxic neurosurgical injections are an important technique in
neuroscience that requires expensive equipment, hardware, and the
development of a high level of skill and precision. There is an urgent
need to reduce the economic and knowledge barrier to making this technique
more accessible. In this work, we have addressed the hardware, software,
and knowledge barriers needed to accomplish successful stereotaxic
intracranial injections in mice using the UD Neuroinjector OSE. First,
the UD Neuroinjector OSE consists of a robust hardware and firmware design
for an automated injector and Arduino-based control system. Using
high resolution image analysis, we have shown stable and highly linear meniscus displacement rates for low volume and low flow-rate injections. Next, we have shown using \textit{in vivo} intracranial
surgery and injections into the mouse brain that the UD Neuroinjector has similar
performance to a commercial injector at less than $0.1\times$ the
cost. Finally, we have designed the UD Neuroinjector Assistant that uses
AI-driven domain expertise via a RAG approach that is grounded in a literature-derived corpus. A CCFv3-based atlas helps guide and complement LLM
responses and target identification assistance. All the components of the
UD Neuroinjector OSE are open source. The UD Neuroinjector is the first open-source end-to-end workflow solution for intracranial injections into the mouse brain.

Automated injection of gene delivery vectors into the brain during
stereotaxic surgery is a superior method compared to manual injection
\cite{Brooks1998CNS}. The approach to design automated injectors has not
significantly changed since it was first systematically demonstrated in
the mouse brain \cite{Brooks1998CNS} indicating design convergence owing
to the constraint of the objective--using a syringe for injection
\cite{Norman2014Innovation}. Many research groups use pressure-based
approaches that use glass micropipettes for injection
\cite{Cetin2006Stereotaxic}. Multiple open-source solutions using and
validating this design and approach have been reported previously
\cite{Yang2011Hunger,Wijnen2014SyringePump,Booeshaghi2019Poseidon,
Arnold2026,Dominguez2026Microinjector}. However, this approach
needs the use of other expensive equipment such as pipette pullers,
manual calibration of volumes, and may often lead to unused injectate due
to calibration errors, which could be relatively expensive depending on
the cost of the vectors. This approach also leads to reduced
standardization across labs. Since syringes used for stereotaxic
injections (such as Hamilton syringes) are easily sterilized, can be
repeatably used for many injections, and are relatively inexpensive
($\sim$100 USD--150 USD per syringe), using this setup can provide
significant cost and time savings for stereotaxic injections.

Commercial automated injector systems rely on proprietary hardware design
and closed-access firmware and software. Beyond the warranty period, the
cost via the primary vendor or even a third-party vendor to maintain and
repair these injector systems following hardware or software malfunction
can be very high. Open-source integrated hardware and software systems can
serve as effective solutions to this problem. Similar to our open-source
design (Figure~1a--c), other recent open-source designs for automated
injectors that use stepper motors have been reported
\cite{Dodson2018SyringePump,BravoMartinez2023Microinjector}. However, head-to-head comparison to commercial injector performance for \textit{in vivo} injections is not available for these designs
\cite{Dodson2018SyringePump,BravoMartinez2023Microinjector}. Our
capillary fluid displacement measurements provide a robust readout of
flow-rate stability using standard operating conditions for stereotaxic
intracranial injections (Fig.~\ref{fig2}). While Bravo-Martinez et al. (2023)
provide a robust design for their stereotaxic injector, they have also
not provided head-to-head comparison to a commercial injector
\cite{BravoMartinez2023Microinjector}. The UD Neuroinjector performs similarly to a leading commercial alternative (QSI, Stoelting Inc.) for \textit{in vivo} performance of stereotaxic injections into the mouse brain
(Fig.~\ref{fig3}).

The open science movement has led to the formation of technology
ecosystems where software, hardware, and user/developer communities
interact. This approach is a welcome change from the siloed development
of scientific technology and has the potential to increase sustainability.
A successful example of this ecosystem is the Bonsai visual programming
platform \cite{Lopes2015Bonsai,Lopes2021Bonsai}. The large user base of
Bonsai programming has contributed to the adoption of Bonsai by commercial
vendors as well \cite{MBFBiosciences2026FP3002}. Our work provides an
integrated framework for the hardware and customizable software
\cite{Sathyanesan2026Configurator} needed for operation of the UD
Neuroinjector. However, we also extend this framework to also provide
domain expertise using a RAG-based AI-driven application (Fig.~\ref{fig4}).
Using our CCFv3-based atlas module and the RAG-based AI assistant tool,
users can easily verify previously reported stereotaxic coordinates into
the specific mouse brain region they are aiming to target. We have
provided validation of AI-assisted target guidance for the cerebellar
simplex (Fig.~\ref{fig4}f) as a proof-of-concept. While it is likely these
extracted coordinates (Fig.~\ref{fig4}f) may not match the actual targets from
the original papers on a one-to-one basis, the fact that these targets
map onto the correct anatomical sub-structure within our atlas module
reference system is noteworthy.

While the UD Neuroinjector possesses many advantages compared to previous open source and commercial solutions, we would like to note a few limitations. We did not opt to design a separate display and input unit for operating the UD Neuroinjector since it reduces build time and users can readily use laptops to perform this function. On the other hand, it would be beneficial to have a standalone display and input unit using low-cost hardware to allow users maximal operational flexibility. While our syringe configurator allows users to customize the control script, our hardware design prevents the use of recent syringe models. We plan to increase the number of compatible syringes in a future release. Our \textit{in vivo} DAPI injections used three mice per injector, however, following flow-rate linearity analysis and \textit{in vivo} validation, the UD Neuroinjector has become our standard injector-of-use for all intracranial stereotaxic injections. For the UD Neuroinjector Assistant, we plan to continue regular expansion of the stereotaxic coordinate database for the mouse brain. Additionally, the absence of rat stereotaxic coordinates represents a significant limitation. Adding a dedicated rat brain stereotaxic coordinate file and atlas module will enable users who perform rat intracranial surgeries access to the UD Neuroinjector Assistant for their workflows. Finally, while our OSE provides individual solutions to each step of the intracranial stereotaxic injection workflow, a more comprehensive and completely integrated system which can use a robotic framework to perform surgeries \cite{rynes2020assembly,coffey2013building} and incorporate AI and agentic capability for human-in-the-loop surgery planning would represent a significant improvement. 

The UD Neuroinjector ecosystem provides a complete end-to-end solution
needed to perform successful literature-validated stereotaxic intracranial
injections into the mouse brain. By democratizing each step of the
workflow, our work brings together a unique combination of tools that will
enable users to significantly reduce the time, effort, and money required
to set up this important and advanced neuroscience workflow in their lab.


\section*{Acknowledgements}

We acknowledge the University of Dayton School of Engineering
Multidisciplinary Design Capstone course sequence (ECE/MEE 431L-432L;
Reilly Downing, Owen Beer, and Megan LaBelle were students in this course)
for providing initial resources for the development of the UD
Neuroinjector. We acknowledge Doug Bishop and Brian LeMaster (AI
Applications and Services, University of Dayton Information Technology)
for technical guidance and providing access to the University of Dayton
FlyerGPT Azure resources. We also gratefully acknowledge the Ohio
Supercomputer Center (OSC) for providing computational resources to Aaron
Sathyanesan (Project \# PNS 0489).


\section*{Author contributions}

Conceptualization: Mir Abbas Raza, Krishna Bhavithavya Kidambi, and Aaron Sathyanesan;
Methodology: Mir Abbas Raza, Reilly Downing, Owen Beer, Megan LaBelle,
Krishna Bhavithavya Kidambi, and Aaron Sathyanesan; Formal analysis: Mir Abbas Raza
and Aaron Sathyanesan; Funding acquisition and Resources: Krishna Bhavithavya Kidambi
and Aaron Sathyanesan; Investigation: Mir Abbas Raza, Sanjay Madala, and
Kassidy Schroeder; Software: Reilly Downing and Aaron Sathyanesan; Project
Administration and Supervision: Krishna Bhavithavya Kidambi and Aaron Sathyanesan;
Visualization: Mir Abbas Raza, Sanjay Madala, Kassidy Schroeder, and Aaron
Sathyanesan; Writing--original draft: Aaron Sathyanesan; Writing--review
and editing: all authors.


\section*{Declarations}

The authors have no relevant financial or non-financial interests to
disclose.
%%%%%%%%%%%%%%%%%%%%%%%%%%%%%%%%%%%%%%%%%%%%%%%%%%%%%%%%%%%%%%%%%%%%
\section*{Supplementary Information}

\begin{description}
  \item[Online Resource 1] (ESM\_1.csv) Spreadsheets containing ground truth values of 84 stereotaxic coordinates from Nucleus accumbens, Hippocampal CA1 region, and Caudate putamen.
  \item[Online Resource 2] (ESM\_2.mmd) Mermaid diagram of the UD Neuroinjector Assistant.
  \item[Online Resource 3] (ESM\_3.mp4) Screen recording of CCFv3-based atlas module and surgery assistant LLM operation.
\end{description}

\section*{Data availability statement}
Hardware design files, firmware, and web application code are publicly available in a Zenodo repository at https://doi.org/10.5281/zenodo.22871234. The experimental datasets generated and analyzed in this study will be made available to peer reviewers on request and will be deposited in a public repository upon acceptance of the manuscript.

%%%%%%%%%%%%%%%%%%%%%%%%%%%%%%%%%%%%%%%%%%%%%%%%%%%%%%%%%%%%%%%%%%%%
\bibliography{sn-bibliography}% common bib file


\begin{figure}[p]
	\centering
    \begin{adjustbox}{max size={\textwidth}{\textheight}}
        \includegraphics{Figure1.png}
    \end{adjustbox}
	\caption{\textbf{UD Neuroinjector Hardware Design and Control.} \textbf{a.} A photograph of the UD Neuroinjector mounted onto the arm of a mouse stereotaxic setup. \textbf{b.} Exploded view of the components of the UD Neuroinjector \textbf{c.} Wiring diagram (fritzing) of Arduino microcontroller-based control of the NEMA 8 motor in the UD Neuroinjector; inset (\textbf{c’}) shows limit switch placement to prevent damage. \textbf{d.} Serial-programming-based interface for controlling the UD Neuroinjector. The script for UD Neuroinjector control and serial programming can be downloaded from the UD Neuroinjector GitHub repository.}
	\label{fig1}
\end{figure} 

\begin{figure}
	\centering
	\includegraphics[width=\linewidth]{Figure2.png}
	\caption{\textbf{Flow rate validation for UD Neuroinjector. a.} Fluid capillary displacement setup showing the black band (‘band’) reference, syringe injection needle, and the water meniscus at the air-water interface inside the capillary. \textbf{b.} Line intensity scan profile of the orange line in panel a showing the inverted peak (orange) corresponding to the reference black band as well as the meniscus profile peak (blue). \textbf{c.} Line scan kymographs for the center of the black reference band (orange) and \textbf{d.} the meniscus edge (blue) for a representative 200 nl/min injection. \textbf{e.} Meniscus relative displacement over the course of the injection for 10 nl/min \textbf{f.} 100 nl/min and \textbf{g.} 200 nl/min flow rate. \textbf{h.} Residual CV comparison between the different flow rate measurements (no statistically significant difference [n.s.] between CV across flow rates; N = 3 injections per group; One-way ANOVA, P = 0.3043). \textbf{i.} Comparison of monotonic fraction across different flow rates (no statistically significant difference [n.s.] between monotonic fraction across flow rates; N = 3 injections per group; One-way ANOVA, P = 0.2071)}
	\label{fig2}
\end{figure}

\begin{figure}
	\centering
	\includegraphics[width=\linewidth]{Figure3.png}
	\caption{\textbf{\textit{In vivo} head-to-head performance of UD Neuroinjector vs. a commercial injector.} \textbf{a.} Representative fluorescence microscopy images of mouse brain tissue sections following stereotaxic surgery and injection of 100 nl of 5 mg/ml DAPI solution using the Quintessential Stereotaxic Injector (QSI, Stoelting Company) (left image) and the UD Neuroinjector (right image) (AP $\sim -1.2$~ to $-1.4$ mm, ML $\sim 0.9$~ to $1.2$ mm, DV $\sim -2.6$~ to $-2.8$ mm); DAPI shown in yellow, anti-NeuN pan-neuronal marker shown in magenta. \textbf{b.} Area of spread comparison between the commercial injector and the UD Neuroinjector shows no statistically significant (n.s.) differences in area of spread (N = 3 mice, n = 2 slices/mouse; Welch’s independent-samples t-test on per-animal mean measurement; Welch’s t = -0.5313, two-sided P = 0.6392). \textbf{c.} No statistically significant differences in DAPI+ cell counts (N = 3 mice, n = 2 slices/mouse; Welch’s independent-samples t-test; Welch’s t = -0.2889, two-sided P = 0.792) between the commercial injector and the UD Neuroinjector. Scale bar for panel a = 150 µm.}
	\label{fig3}
\end{figure}

\begin{figure}
	\centering
	\includegraphics[width=\linewidth]{Figure4.png}
	\caption{\textbf{Corpus-grounded AI-driven domain expertise and knowledge layer for the UD Neuroinjector ecosystem.} \textbf{a.} Initial scoping of PubMed Central (PMC) for search terms associated with injection, surgery, and mouse brain regions yielded an initial corpus of 18,283 papers. \textbf{b.} We used GPU-accelerated distillation of the papers using the \texttt{gpt-oss-20b} open source model on the Ohio Supercomputer and obtained \textbf{c.} a structured file, \texttt{injection-structured.json}. \textbf{d.} The \texttt{injection-structured.json} file contains 3,738 PMIDs with at least one complete (antero-posterior – AP, mediolateral – ML, and dorsoventral – DV) coordinate set representing 26.6\% of the 14,057 PMIDs. \textbf{e.} An analysis of the targets per common coordinate framework (CCF) division (n = 9,184 targets assigned) reveals the spread of the coordinates, with the hippocampus having the largest number of previously reported coordinates followed by the striatum. \textbf{f.} Proof-of-feasibility mapping of extracted coordinates (cyan circles) onto the CCFv3 coordinate system for the cerebellar simplex (magenta) found in PMIDs 31934857 (AP -5.80 relative to bregma), 32795386 (AP -6.10 relative to bregma), 38605812 (AP -6.20; relative to bregma but rederived from lambda – 4.20), and 40848722 (AP -7.00; relative to bregma but rederived from lambda – 4.20).}
	\label{fig4}
\end{figure}


\end{document}
