## Supplementary material for "An integrated stereotaxic injection architecture for mouse intracranial surgeries with AI-driven domain expertise": Manuscript LaTeX Project Zip: sn-bibliography.bib

@article{coffey2013building,
  author    = {Coffey, K. R. and Barker, D. J. and Ma, S. and West, M. O.},
  title     = {Building an open-source robotic stereotaxic instrument},
  journal   = {Journal of Visualized Experiments},
  number    = {80},
  pages     = {e51006},
  year      = {2013},
  doi       = {10.3791/51006},
  pmid      = {24192514},
  pmcid     = {PMC3965336}
}

@article{rynes2020assembly,
  title={Assembly and operation of an open-source, computer numerical controlled (CNC) robot for performing cranial microsurgical procedures},
  author={Rynes, Mathew L and Ghanbari, Leila and Schulman, Daniel Sousa and Linn, Samantha and Laroque, Michael and Dominguez, Judith and Navabi, Zahra S and Sherman, Peter and Kodandaramaiah, Suhasa B},
  journal={Nature protocols},
  volume={15},
  number={6},
  pages={1992--2023},
  year={2020},
  publisher={Nature Publishing Group UK London}
}

@article{comeau2019pmc,
  title={PMC text mining subset in BioC: about three million full-text articles and growing},
  author={Comeau, Donald C and Wei, Chih-Hsuan and Islamaj Do{\u{g}}an, Rezarta and Lu, Zhiyong},
  journal={Bioinformatics},
  volume={35},
  number={18},
  pages={3533--3535},
  year={2019},
  publisher={Oxford University Press}
}

@incollection{sayers2008quickstart,
  author       = {Sayers, Eric},
  title        = {{E}-utilities {Q}uick {S}tart},
  booktitle    = {Entrez {P}rogramming {U}tilities {H}elp [{I}nternet]},
  year         = {2008},
  publisher    = {National Center for Biotechnology Information (US)},
  address      = {Bethesda (MD)},
  url          = {https://www.ncbi.nlm.nih.gov/books/NBK25500/},
  note         = {Available from: \url{https://www.ncbi.nlm.nih.gov/books/NBK25500/}}
}

@article{claudi2021visualizing,
  title={Visualizing anatomically registered data with brainrender},
  author={Claudi, Federico and Tyson, Adam L and Petrucco, Luigi and Margrie, Troy W and Portugues, Ruben and Branco, Tiago},
  journal={Elife},
  volume={10},
  pages={e65751},
  year={2021},
  publisher={eLife Sciences Publications, Ltd}
}

@article{fuglstad2023histological,
  title={Histological E-data registration in rodent brain spaces},
  author={Fuglstad, Jingyi Guo and Saldanha, Pearl and Paglia, Jacopo and Whitlock, Jonathan R},
  journal={Elife},
  volume={12},
  pages={e83496},
  year={2023},
  publisher={eLife Sciences Publications, Ltd}
}

@book{franklin2019paxinos,
  title={Paxinos and Franklin's the Mouse brain in stereotaxic coordinates, compact: The coronal plates and diagrams},
  author={Franklin, Keith BJ and Paxinos, George},
  year={2019},
  address={San Diego},
  publisher={Academic Press}
}

@article{baden2015open,
  title={Open Labware: 3-D printing your own lab equipment},
  author={Baden, Tom and Chagas, Andre Maia and Gage, Greg and Marzullo, Timothy and Prieto-Godino, Lucia L and Euler, Thomas},
  journal={PLoS biology},
  volume={13},
  number={3},
  pages={e1002086},
  year={2015},
  publisher={Public Library of Science San Francisco, CA USA}
}

@article{tye2012optogenetic,
  title={Optogenetic investigation of neural circuits underlying brain disease in animal models},
  author={Tye, Kay M and Deisseroth, Karl},
  journal={Nature Reviews Neuroscience},
  volume={13},
  number={4},
  pages={251--266},
  year={2012},
  publisher={Nature Publishing Group UK London}
}

@article{Arnold2026,
	author = "Jon Arnold and Jason Osborne",
	title = "{Stereotaxic Injector System}",
	year = "2026",
	month = "4",
	url = "https://janelia.figshare.com/articles/physical_object/Stereotaxix_Injector_System/31955799",
	doi = "10.25378/janelia.31955799.v2"
}

@article{Berg2019Ilastik,
	title={Ilastik: interactive machine learning for (bio) image analysis},
	author={Berg, Stuart and Kutra, Dominik and Kroeger, Thorben and Straehle, Christoph N and Kausler, Bernhard X and Haubold, Carsten and Schiegg, Martin and Ales, Janez and Beier, Thorsten and Rudy, Markus and others},
	journal={Nature methods},
	volume={16},
	number={12},
	pages={1226--1232},
	year={2019},
	publisher={Nature Publishing Group US New York}
}

@article{Booeshaghi2019Poseidon,
	title={Principles of open source bioinstrumentation applied to the poseidon syringe pump system},
	author={Booeshaghi, A Sina and Beltrame, Eduardo da Veiga and Bannon, Dylan and Gehring, Jase and Pachter, Lior},
	journal={Scientific reports},
	volume={9},
	number={1},
	pages={12385},
	year={2019},
	publisher={Nature Publishing Group UK London}
}

@article{BravoMartinez2023Microinjector,
	author  = {Bravo-Mart{\'i}nez, Jorge and Ortega-Tinoco, Sonia and
	Gardu{\~n}o, Julieta and Hern{\'a}ndez-L{\'o}pez, Salvador},
	title   = {Arduino based intra-cerebral microinjector device for
	neuroscience research},
	journal = {HardwareX},
	year    = {2023},
	volume  = {15},
	pages   = {e00446},
	doi     = {10.1016/j.ohx.2023.e00446}
}

@article{Brooks1998CNS,
	author  = {Brooks, Andrew I. and Halterman, Marc W. and
	Chadwick, Christopher A. and Davidson, Beverly L. and
	Haak-Frendscho, Mary and Radel, Clyde and Porter, Chris and
	Federoff, Howard J.},
	title   = {Reproducible and efficient murine {CNS} gene delivery using
	a microprocessor-controlled injector},
	journal = {Journal of Neuroscience Methods},
	year    = {1998},
	volume  = {80},
	number  = {2},
	pages   = {137--147},
	doi     = {10.1016/S0165-0270(97)00207-0}
}

@article{Cetin2006Stereotaxic,
	author  = {Cetin, Ali and Komai, Shinichiro and Eliava, Marina and
	Seeburg, Peter H. and Osten, Pavel},
	title   = {Stereotaxic gene delivery in the rodent brain},
	journal = {Nature Protocols},
	year    = {2006},
	volume  = {1},
	pages   = {3166--3173},
	doi     = {10.1038/nprot.2006.450}
}

@misc{Dodson2018SyringePump,
	author = {Dodson, Paul},
	title  = {Syringe Pump},
	year   = {2018},
	note   = {Open-source syringe pump}
}

@article{Dominguez2026Microinjector,
	author  = {Dominguez, Victor H. and Frankfurter, Maxwell and
	Hayes, Kevin B. and Kahn, Mark L. and Dominguez, Martin H.},
	title   = {A portable, ultra-low cost, open-source, pedal-controlled
	microinjector for laboratory use},
	journal = {PLOS ONE},
	year    = {2026},
	volume  = {21},
	number  = {5},
	pages   = {e0347487},
	doi     = {10.1371/journal.pone.0347487}
}

@article{Lopes2015Bonsai,
	author  = {Lopes, Gon{\c{c}}alo and Bonacchi, Niccol{\`o} and
	Fraz{\~a}o, Jo{\~a}o and Neto, Joana P. and
	Atallah, Bassam V. and Soares, Sofia and Moreira, Lu{\'i}s and
	Matias, Sara and Itskov, Pavel M. and Correia, Patr{\'i}cia A.
	and Medina, Roberto E. and Calcaterra, Lorenza and
	Dreosti, Elena and Paton, Joseph J. and Kampff, Adam R.},
	title   = {Bonsai: an event-based framework for processing and
	controlling data streams},
	journal = {Frontiers in Neuroinformatics},
	year    = {2015},
	volume  = {9},
	pages   = {7},
	doi     = {10.3389/fninf.2015.00007}
}

@article{Lopes2021Bonsai,
	author  = {Lopes, Gon{\c{c}}alo and Monteiro, Patricia},
	title   = {New Open-Source Tools: Using Bonsai for Behavioral Tracking
	and Closed-Loop Experiments},
	journal = {Frontiers in Behavioral Neuroscience},
	year    = {2021},
	volume  = {15},
	pages   = {647640},
	doi     = {10.3389/fnbeh.2021.647640}
}

@misc{MBFBiosciences2026FP3002,
	author       = {{MBF Biosciences}},
	title        = {{NeuroPhotoMetrics FP3002}},
	year         = {2026},
	url          = {https://web.archive.org/web/20260628175230/https://www.mbfbioscience.com/products/fp3002},
	note         = {Accessed 16 September 2026}
}

@misc{McCauley2026AccelStepper,
	author = {McCauley, Mike},
	title  = {{AccelStepper}: AccelStepper library for Arduino},
	year   = {2026},
	note   = {Arduino stepper motor library}
}

@article{Norman2014Innovation,
	author  = {Norman, Donald A. and Verganti, Roberto},
	title   = {Incremental and Radical Innovation: Design Research vs.
	Technology and Meaning Change},
	journal = {Design Issues},
	year    = {2014},
	volume  = {30},
	number  = {1},
	pages   = {78--96},
	doi     = {10.1162/DESI_a_00250}
}

@misc{OpenAI2025GPTOSS,
	author        = {{OpenAI}},
	title         = {{gpt-oss-120b \& gpt-oss-20b Model Card}},
	year          = {2025},
	eprint        = {2508.10925},
	archivePrefix = {arXiv},
	primaryClass  = {cs.CL},
	url           = {https://arxiv.org/abs/2508.10925}
}

@misc{Sathyanesan2026Assistant,
	author = {Sathyanesan, Aaron},
	title  = {{UD Neuroinjector Assistant}},
	year   = {2026},
	url    = {https://asathyanesan.github.io/Neuroinjector-OSE/assistant/},
	note   = {Accessed 9 September 2026}
}

@software{Raza_UD_Neuroinjector_Open_2026,
author = {Raza, Mir Abbas and Downing, Reilly and Beer, Owen and LaBelle, Megan and Madala, Sanjay and Schroeder, Kassidy and Kidambi, Krishna B. and Sathyanesan, Aaron},
doi = {10.5281/zenodo.22871234},
license = {MIT},
month = sep,
title = {{UD Neuroinjector Open Source Ecosystem}},
url = {https://github.com/asathyanesan/Neuroinjector-OSE},
version = {0.1.0},
year = {2026}
}

@misc{Sathyanesan2026Configurator,
	author = {Sathyanesan, Aaron},
	title  = {{UD Neuroinjector --- Syringe Configurator}},
	year   = {2026},
	url    = {https://asathyanesan.github.io/Neuroinjector-OSE/webapp/},
	note   = {Accessed 12 September 2026}
}

@article{Schindelin2012Fiji,
	author  = {Schindelin, Johannes and Arganda-Carreras, Ignacio and
	Frise, Erwin and Kaynig, Verena and Longair, Mark and
	Pietzsch, Tobias and Preibisch, Stephan and Rueden, Curtis and
	Saalfeld, Stephan and Schmid, Benjamin and Tinevez,
	Jean-Yves and White, Daniel James and Hartenstein, Volker and
	Eliceiri, Kevin and Tomancak, Pavel and Cardona, Albert},
	title   = {Fiji: an open-source platform for biological-image analysis},
	journal = {Nature Methods},
	year    = {2012},
	volume  = {9},
	pages   = {676--682},
	doi     = {10.1038/nmeth.2019}
}

@article{Wang2020Allen,
	author  = {Wang, Quanxin and Ding, Song-Lin and Li, Yang and
	Royall, Joshua and Feng, David and Lesnar, Phil and
	Graddis, Nile and Naeemi, Maitham and Facer, Benjamin and
	Ho, Anh and Dolbeare, Tim and Blanchard, Ben and
	Dee, Nick and Wakeman, Wayne and Hirokawa, Karla E. and
	Szafer, Aaron and Sunkin, Susan M. and Oh, Seung Wook and
	Bernard, Amy and Phillips, John W. and Hawrylycz, Michael and
	Koch, Christof and Zeng, Hongkui and Harris, Julie A. and
	Ng, Lydia},
	title   = {The Allen Mouse Brain Common Coordinate Framework:
	A 3D Reference Atlas},
	journal = {Cell},
	year    = {2020},
	volume  = {181},
	number  = {4},
	pages   = {936--953.e20},
	doi     = {10.1016/j.cell.2020.04.007}
}

@article{Wijnen2014SyringePump,
	author  = {Wijnen, Bas and Hunt, Emily J. and Anzalone, Gerald C. and
	Pearce, Joshua M.},
	title   = {Open-Source Syringe Pump Library},
	journal = {PLOS ONE},
	year    = {2014},
	volume  = {9},
	number  = {9},
	pages   = {e107216},
	doi     = {10.1371/journal.pone.0107216}
}

@article{Yang2011Hunger,
	author  = {Yang, Yunlei and Atasoy, Deniz and Su, Helen H. and
	Sternson, Scott M.},
	title   = {Hunger states switch a flip-flop memory circuit via a
	synaptic {AMPK}-dependent positive feedback loop},
	journal = {Cell},
	year    = {2011},
	volume  = {146},
	number  = {6},
	pages   = {992--1003},
	doi     = {10.1016/j.cell.2011.07.039}
}
