## Supplementary figures and images for "An integrated stereotaxic injection architecture for mouse intracranial surgeries with AI-driven domain expertise"

### Figure1.png

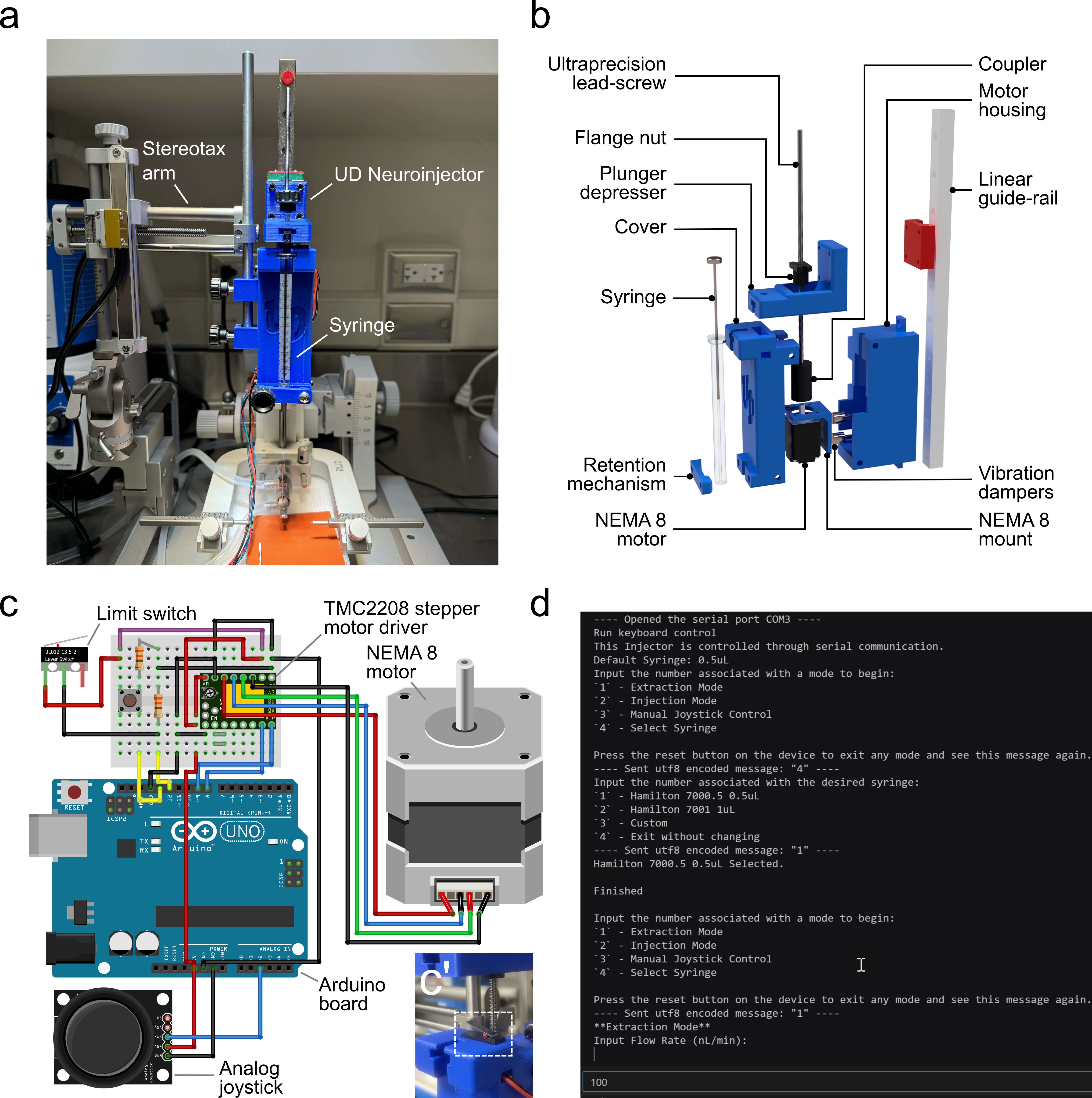

### Figure2.png

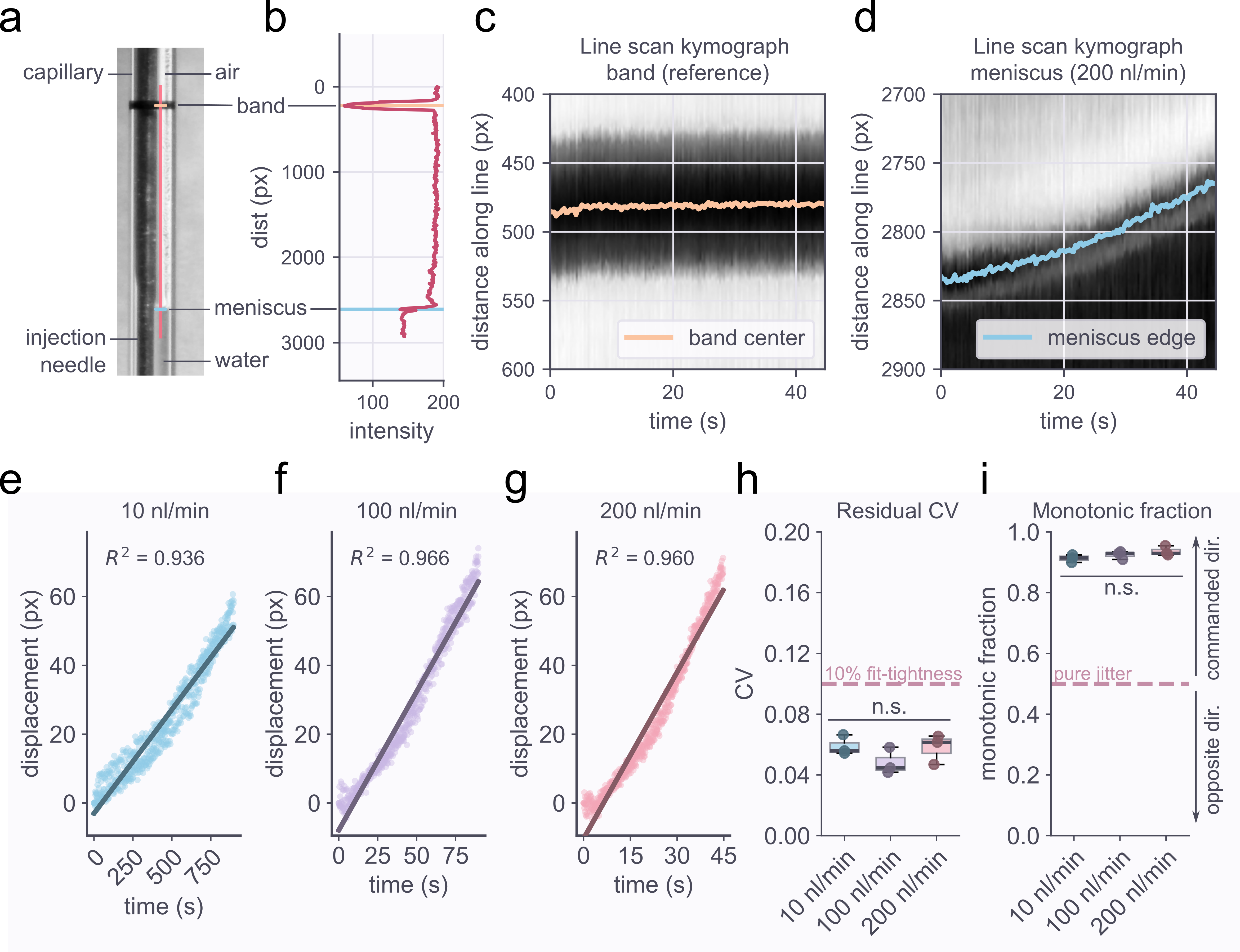

### Figure3.png

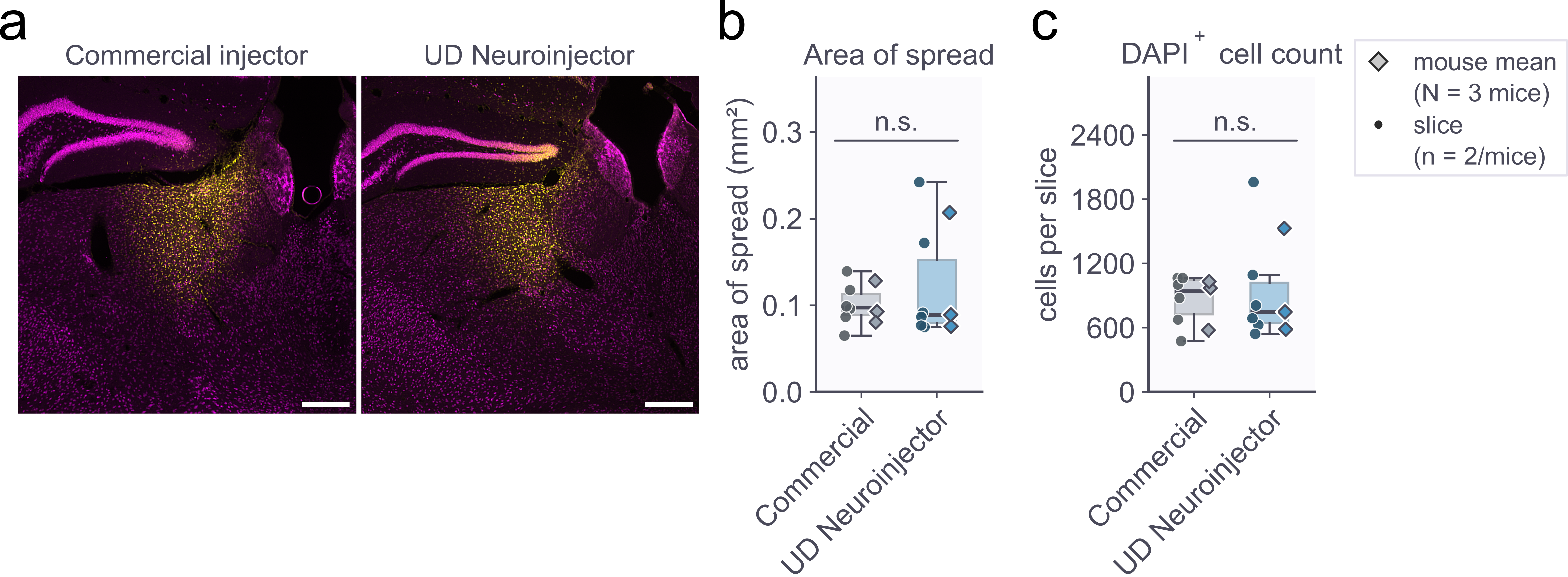

### Figure4.png

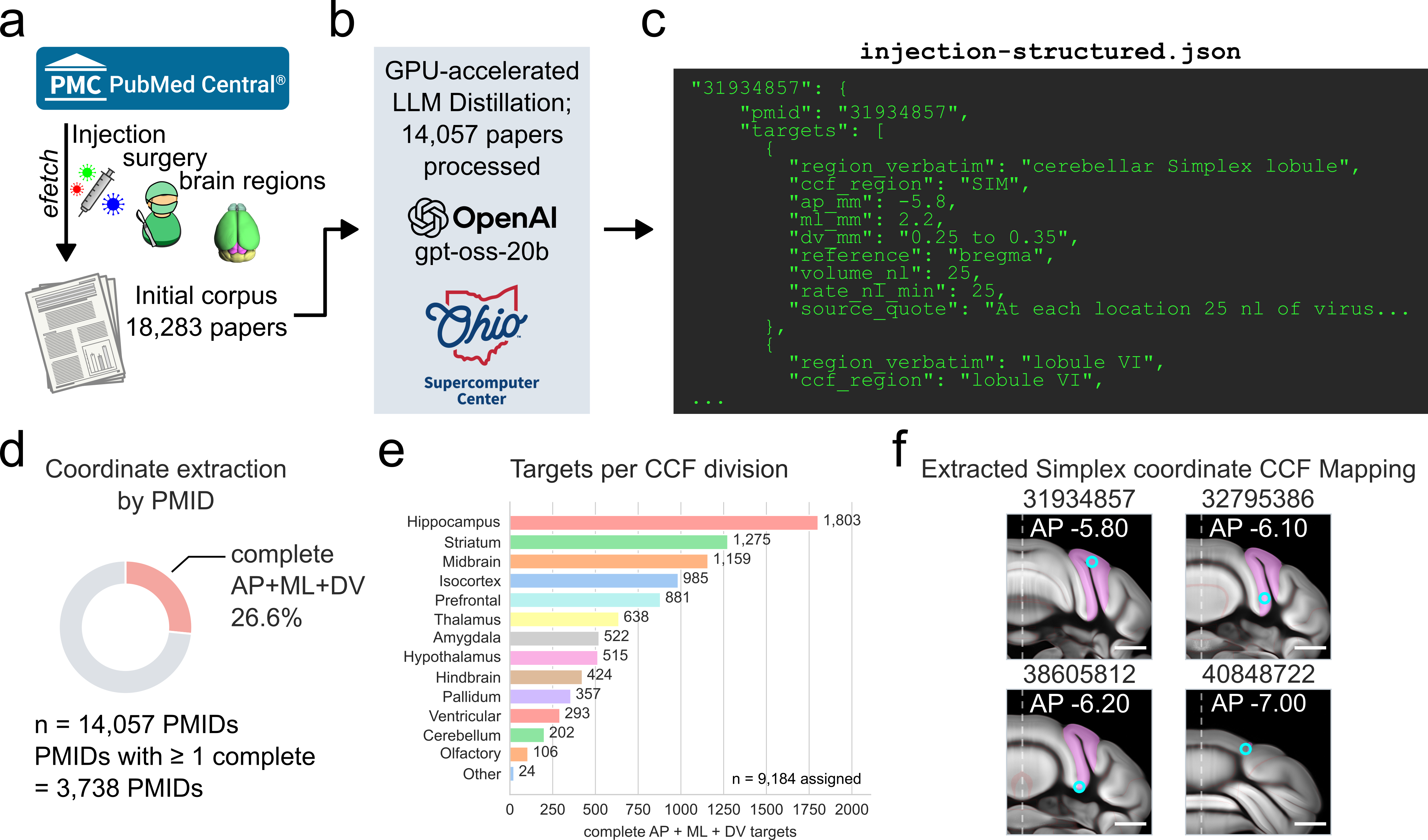
